# Deep mutational scanning reveals functional constraints and antigenic variability of Lassa virus glycoprotein complex

**DOI:** 10.1101/2024.02.05.579020

**Authors:** Caleb R. Carr, Katharine H. D. Crawford, Michael Murphy, Jared G. Galloway, Hugh K. Haddox, Frederick A. Matsen, Kristian G. Andersen, Neil P. King, Jesse D. Bloom

## Abstract

Lassa virus is estimated to cause thousands of human deaths per year, primarily due to spillovers from its natural host, *Mastomys* rodents. Efforts to create vaccines and antibody therapeutics must account for the evolutionary variability of Lassa virus’s glycoprotein complex (GPC), which mediates viral entry into cells and is the target of neutralizing antibodies. To map the evolutionary space accessible to GPC, we use pseudovirus deep mutational scanning to measure how nearly all GPC amino-acid mutations affect cell entry and antibody neutralization. Our experiments define functional constraints throughout GPC. We quantify how GPC mutations affect neutralization by a panel of monoclonal antibodies and show that all antibodies are escaped by mutations that exist among natural Lassa virus lineages. Overall, our work describes a biosafety-level-2 method to elucidate the mutational space accessible to GPC and shows how prospective characterization of antigenic variation could aid design of therapeutics and vaccines.

## Introduction

Lassa virus is the causative agent of Lassa fever, which has a high fatality rate in hospitalized cases and causes thousands of deaths each year across West Africa.^1–6^ Lassa virus is almost exclusively transmitted to humans from spillovers from the natural reservoir host *Mastomys natalensis,*^5,7,8^ but human-to-human transmission has been documented, especially during nosocomial outbreaks.^9,10^ Currently, there are no approved vaccines or therapeutics except for off-label use of ribavirin, which has debatable benefits.^11–14^ The development of vaccines and therapeutics is further complicated by the high level of genetic diversity of Lassa virus, with seven designated lineages occupying distinct geographical locations.^10,15–19^

An important target for vaccines and therapeutics is the glycoprotein complex (GPC), the only viral surface protein. GPC is expressed as a polypeptide precursor that is cleaved by signal peptidase^20,21^ and subtilisin kexin isozyme-1 / site-1 protease^22–25^ resulting in three distinct polypeptides that remain physically associated: stable signal peptide (SSP), glycoprotein 1 (GP1), and glycoprotein 2 (GP2). The mature GPC exists on the viral surface as a homotrimer and mediates viral entry via the primary host receptor *ɑ*-dystroglycan (*ɑ*-DG)^26^ and secondary host receptors, such as LAMP1.^27^

Anti-GPC antibodies that can neutralize Lassa virus have been isolated from Lassa fever survivors.^28^ Some of these antibodies are being developed as antibody cocktails and there is evidence that they can protect from lethal Lassa virus infection.^29–32^ In addition, vaccines that elicit antibodies against GPC are also being developed.^33–37^ However, the high variability of GPC (up to ∼8% amino-acid divergence among lineages) raises the specter of antibody and vaccine escape, a problem that has been observed for other viruses—including most recently SARS-CoV-2.^38–41^

To understand the potential for genetic and antigenic variation in GPC, we used deep mutational scanning^42^ to measure how nearly all GPC amino-acid mutations affect cell entry and antibody neutralization in the context of a safe pseudovirus system. Briefly, these experiments involve making large libraries of GPC mutants, and then measuring the effects of mutations in pooled infection experiments followed by deep sequencing. The resulting maps of mutational effects highlight mutationally tolerant and intolerant regions of GPC, and identify specific mutations that escape antibody neutralization. These maps can inform the development of therapeutic antibodies and vaccines with increased robustness to Lassa virus antigenic variation. To help aid these efforts, we provide interactive visualizations of our GPC maps, enabling end-users to explore and investigate the functional and antigenic effects of individual Lassa virus mutations.

## Results

### Pseudovirus deep mutational scanning of GPC

To study the impacts of mutations to the Lassa virus GPC, we utilized a pseudovirus deep mutational scanning platform that we previously developed and applied to HIV envelope^43^ and SARS-CoV-2 spike.^42,44^ This platform enabled us to create large libraries of GPC-pseudotyped lentiviruses that have a genotype-phenotype link between a barcode in the lentivirus genomes and the GPC variant on the surface of each virion (Figure 1A,B). Because the GPC-pseudotyped lentiviruses can only undergo a single round of infection and encode no viral genes other than GPC, they are safe to study at biosafety-level-2^45^ even though Lassa virus itself is a biosafety-level-4 agent.^46^

**Figure 1.**
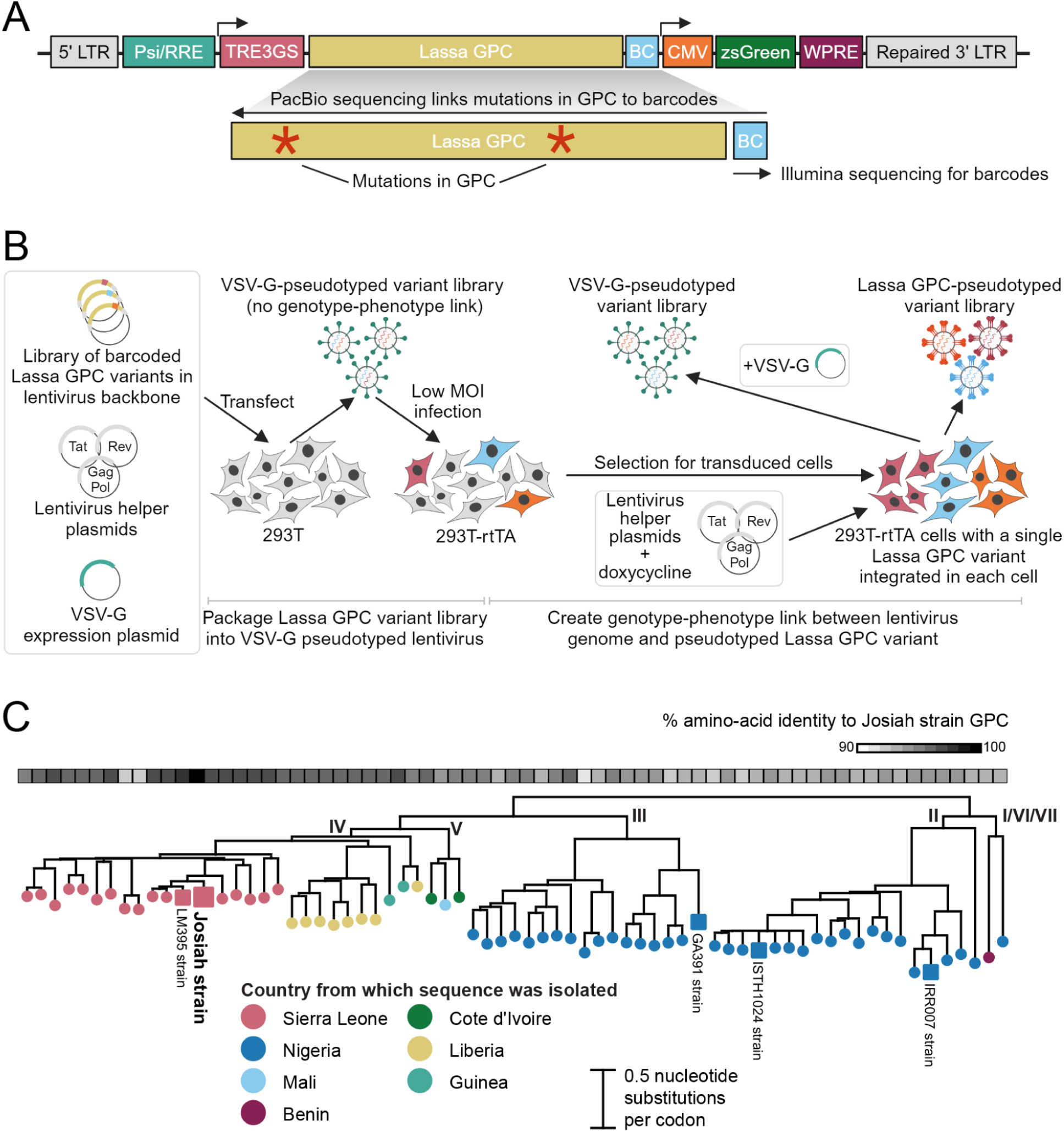
Pseudovirus deep mutational scanning of Lassa GPC. **A** Lentivirus backbone used for GPC deep mutational scanning. The backbone contains full-length 5′ and 3′ long terminal repeat (LTR) sequences. GPC is under control of a doxycycline-inducible TRE3GS promoter and linked to a random 16-nucleotide barcode (BC) downstream of the stop codon. A CMV promoter drives the expression of zsGreen. Other backbone components include the lentiviral Psi/Rev response element (RRE), woodchuck hepatitis virus post-transcriptional regulatory element (WPRE), and a repaired 3’-LTR to allow re-activation of integrated genomes.^42,92^ **B** Approach for producing genotype-phenotype linked GPC-pseudotyped lentivirus libraries. GPC-encoding lentivirus backbone, vesicular stomatitis virus G protein (VSV-G) expression plasmid, and lentivirus helper plasmids are first transfected into 293T cells to produce a VSV-G-pseudotyped variant library with no genotype-phenotype link. To create a genotype-phenotype link, the VSV-G-pseudotyped variant library is used to infect reverse tetracycline-controlled transactivator (rtTA) expressing 293T cells at low multiplicity of infection (MOI) so infected cells typically are transduced with just one lentivirus genome. Transduced cells are selected based on expression of zsGreen. Finally, GPC mutant viruses with a genotype-phenotype link are generated from the transduced cells by inducing GPC expression with doxycycline and transfecting the lentivirus helper plasmids. The library composition can be assessed by separately transfecting the transduced cells with VSV-G alongside the helper plasmids, which creates VSV-G-pseudotyped viruses that infect cells regardless of the functionality of the GPC variant encoded in the lentivirus genome. **C** Phylogenetic tree of representative Lassa GPC sequences. Tree tips are colored by the country from which the virus was collected. The major Lassa lineages I, II, III, IV, V, VI, and VII are labeled. The GPC from the Josiah strain used for deep mutational scanning is labeled in large bold font. Other GPC sequences used later in this paper are labeled in smaller font. Percent amino-acid identity with respect to the Josiah strain GPC is shown for all sequences above the tree.

For our deep mutational scanning experiments, we used GPC from the Lassa virus lineage IV Josiah strain (Figure 1C).^17,47^ We chose the Josiah strain because it is widely used in studies of candidate Lassa virus therapeutics and vaccines.^27,28,30,31,48–52^ Importantly, we could obtain high titers of lentiviruses pseudotyped with this GPC in our deep mutational scanning platform (Figure S1A). The Josiah GPC differs by ∼8% at the protein level (∼40 amino-acid mutations) from the GPCs of the most distantly related Lassa virus lineages (Figure 1C).

We created deep mutational scanning libraries using a PCR-based mutagenesis method that introduces all amino acid mutations at each site in GPC.^42,43,53,54^ The GPC variants in our libraries contained an average ∼2 nonsynonymous mutations each (Figure S1B), with the number of amino-acid mutations per variant roughly following a Poisson distribution (Figure S1C). We generated two independent GPC mutant libraries each containing ∼50,000 variants that covered ∼99% of all 9,820 possible amino-acid mutations to GPC (Figure S1D).

### Effects of mutations on GPC-mediated viral entry

Because Lassa virus normally circulates in its reservoir host of *Mastomys* (mastomys) rodents, its GPC should be optimized by evolution for efficient infection of these species. However, human Lassa virus infections typically represent spillovers,^7,17,55^ so it is conceivable that GPC may not be well adapted to infect humans. Because Lassa virus is an agent with potential biosecurity relevance,^56^ we were cognizant that our study (which uses pseudovirus experiments that are themselves safe) could generate potentially hazardous information^57,58^ about mutations that specifically adapt GPC to human cell entry.^26,27,59^ GPC binds to a specific glycan (matriglycan) on its primary receptor *ɑ*-DG. This glycosylation site is conserved between human and mastomys *ɑ*-DG orthologs (Figure S2).^59–63^ We therefore anticipated that the effects of mutations on GPC-mediated entry were likely to be similar regardless of whether the target cells expressed human or mastomys *ɑ*-DG—but since this assumption has never been experimentally tested, we first set out to do that. We decided that if we identified mutations that specifically adapted GPC to better enter cells expressing human *ɑ*-DG, then we would consider reporting just results from experiments using cells expressing only mastomys *ɑ*-DG to avoid providing information^57,58^ about specific adaptations for infection of cells expressing human *ɑ*-DG.

To screen for GPC mutations that improved use of human versus mastomys *ɑ*-DG, we first created stable cell lines that express either human or mastomys *ɑ*-DG by introducing the *DAG1* gene that encodes *ɑ*-DG into 293T cells lacking that gene (293TΔDAG1 cells^27^) (Figure S3). Next, we determined the effects of GPC mutations on cell entry^42^ for 293T, 293TΔDAG1+humanDAG1, and 293TΔDAG1+mastomysDAG1 cell lines by infecting cells with either GPC-pseudotyped or VSV-G-pseudotyped variant libraries and sequencing the variant barcodes from the infected cells (Figure S4). All variants infect cells when VSV-G is present, but only variants with functional GPC infect cells when VSV-G is not present. The enrichment or depletion of variants in the GPC-pseudotyped condition relative to the VSV-G-pseudotyped condition quantifies their capability to enter cells. The effects of mutations on entry into 293T, 293TΔDAG1+humanDAG1, and 293TΔDAG1+mastomysDAG1 cells were highly correlated (Figure S5A), and we identified no mutations that specifically improved use of human versus mastomys *ɑ*-DG. This finding was confirmed by further analysis of the deep mutational scanning data by an algorithm^64^ that looked for statistical evidence of mutations with different effects on entry into cells expressing human versus mastomys *ɑ*-DG (Figure S5B-D). Note that our experiments do not assess other cell-type specific factors, such as variation in the glycosyltransferase responsible for glycosylating *ɑ*-DG^61,65^ or other secondary receptors,^27,66–69^ that could affect GPC-mediated entry into different cells. They also obviously do not assess non-GPC determinants of Lassa virus infection of cells from different species.

Since we did not detect any GPC mutations that differentially affected entry into cells expressing human versus mastomys *ɑ*-DG, for all further experiments we used wild-type 293T cells, which naturally express human *ɑ*-DG. We performed four replicate deep mutational scanning experiments to measure the effects of mutations on entry into 293T cells for each of the two GPC libraries, following the workflow outlined in Figure S4. We quantified a cell-entry score for each barcoded GPC variant, with scores of zero indicating parental-like (wild-type Josiah) entry, negative scores indicating impaired entry, and positive scores indicating improved entry. As expected, unmutated GPC variants and variants with synonymous mutations have cell-entry scores close to zero, while variants with premature stop codons have highly negative scores (Figure S6A). Variants with one nonsynonymous mutation had scores spanning a range from highly negative to slightly positive, whereas variants with multiple nonsynonymous mutations tended to have negative scores due to the accumulation of typically deleterious mutations (Figure S6A).

To estimate the effects of individual GPC mutations on cell entry, we decomposed the entry scores for all single- and multi-mutant variants using global epistasis models (Figure 2A and interactive heatmap at https://dms-vep.github.io/LASV_Josiah_GP_DMS/htmls/293T_entryfunc_effects.html).^64,70^ Inspection of the full heatmap of mutational effects shows many mutations are deleterious to cell entry, but some are well tolerated. Mutational effects on cell entry differ across GPC regions (Figure 2B,C): the stalk (which includes the SSP and part of GP2) is less mutationally tolerant than the head of the ectodomain (which includes most of GP1). The core binding pocket for *ɑ*-DG^52^ is also highly mutationally intolerant; however, a peripheral site, K125, is quite mutationally tolerant, at least in the context of GPC-pseudotyped lentivirus (Figure 2A-C, S6B, and S6C). The residues important for LAMP1^71,72^ binding are often quite tolerant of mutations except for the histidine triad (H92, H93, and H230) (Figure 2A-C, S6B, and S6C). The tolerance of GPC sites to mutations with respect to cell entry as measured by our deep mutational scanning correlates with the observed natural sequence diversity at each site (Figures S6D,E).

**Figure 2.**
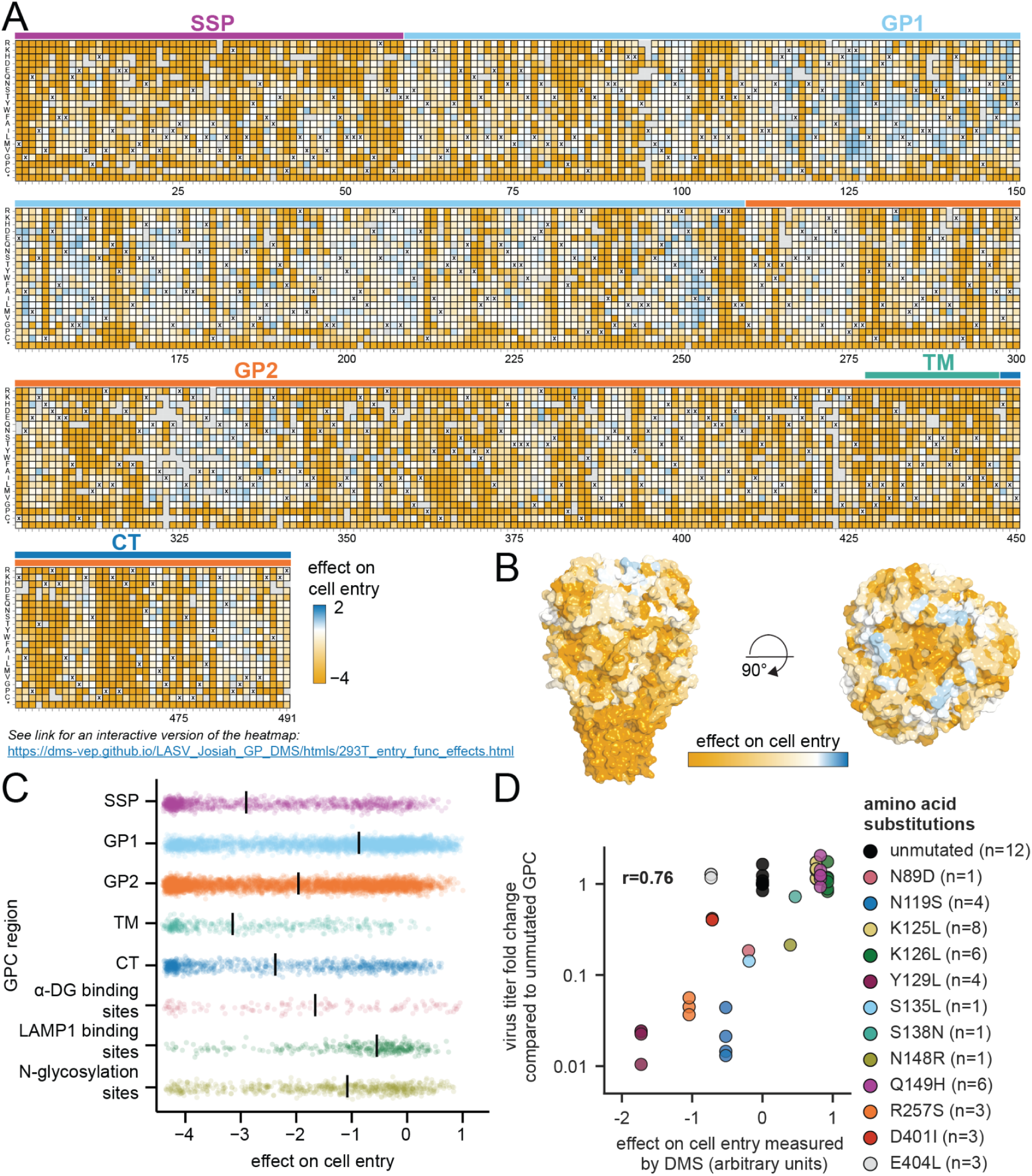
Effects of GPC mutations on cell entry. **A** Effects of mutations on entry into 293T cells as measured by deep mutational scanning. Each square in the heatmap represents a different mutation, with mutations that impair cell entry colored orange and those that improve cell entry colored blue. The wildtype amino-acid in the parental Josiah strain at each site is indicated with a x. The overlay bar denotes the stable signal peptide (SSP), glycoprotein 1 (GP1) domain, glycoprotein 2 (GP2) domain, transmembrane domain (TM), and cytoplasmic tail (CT).^93^ See the interactive version of the heatmap at https://dms-vep.github.io/LASV_Josiah_GP_DMS/htmls/293T_entry_func_effects.html for more effective visualization of the data. **B** Surface representation of GPC colored by the average effect of all amino-acid mutations at each site on cell entry (PDB: 7PUY). **C** Effects of mutations on cell entry for different GPC regions. Each point represents a different mutation, and medians are shown for each region as vertical lines. **D** Correlation of effects on cell entry measured by deep mutational scanning and the fold-change in titer of individual GPC pseudovirus mutants relative to unmutated Josiah GPC. Each point represents a biological replicate. The number of biological replicates is indicated in the legend for each mutant. The Pearson correlation (r) is indicated.

We validated the deep mutational scanning cell-entry measurements for specific mutations with single mutant GPC-pseudotyped lentiviruses (Figure 2D).^73^ The infectious titers of single-mutant GPC pseudoviruses correlated with the deep mutational scanning measurements (Figure 2D).

### Mapping of GPC mutations that escape antibody neutralization

Antibodies are being developed as therapeutics and prophylactics for Lassa virus,^29,30,32,74^ but pre-existing genetic variation or ongoing evolution can render antiviral antibodies ineffective.^38–41^ To assess the potential for GPC to acquire antibody-escape mutations, we used deep mutational scanning to measure how GPC mutations affect antibody neutralization. To do this, we incubated the pseudovirus libraries with increasing concentrations of antibody before infecting target cells, using a non-neutralized VSV-G-pseudotyped “standard” virus to convert sequencing counts to absolute neutralization values (Figure 3A).^42,43^ We then analyzed the data using a biophysical model^75^ that computes an escape value for each mutation that is roughly proportional to the log fold change in antibody inhibitory concentration 50% (IC_50_) caused by the mutation.

**Figure 3.**
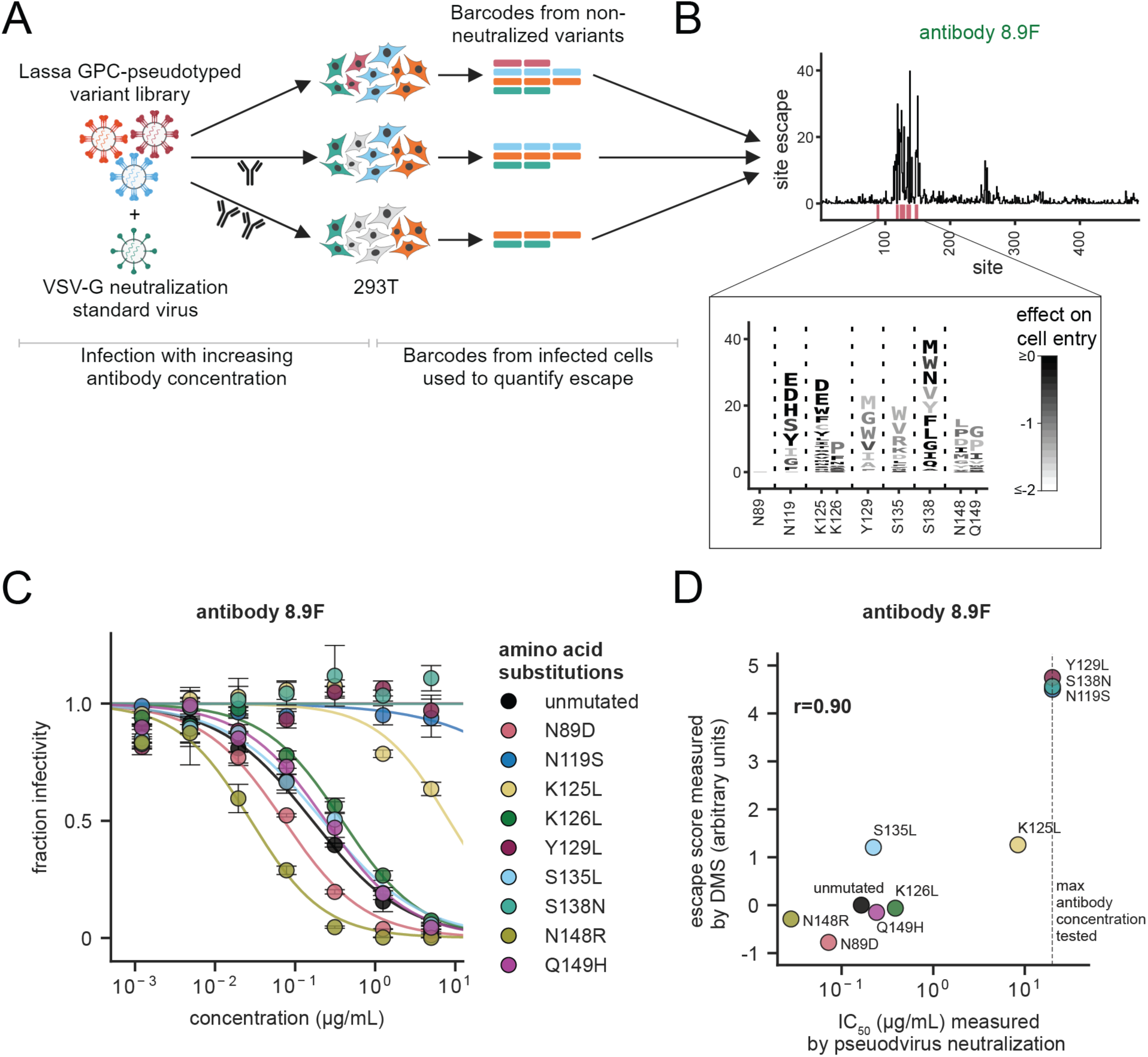
Mapping the effects of mutations on antibody escape. **A** Workflow for antibody-escape mapping. The GPC pseudovirus library is mixed with a VSV-G-pseudotyped “standard” that is not neutralized by anti-Lassa antibodies. The pseudovirus pool is incubated with different antibody concentrations, then used to infect cells. Viral genomes are recovered from infected cells ∼12 hours post infection, and barcodes are sequenced. Sequencing counts are normalized to the VSV-G standard to compute neutralization. **B** Escape from the antibody 8.9F visualized as a line plot showing summed effects of all escape mutations at each site, or a logo plot where the height of each letter indicates the escape caused by that mutation. Letters are colored by effects of mutations on cell entry in the absence of antibody, with mutations that impair entry in lighter gray. The sites of mutations chosen for validation as described in **C** and **D** are shown in the logo plot and highlighted pink below the line plot. See https://dms-vep.org/LASV_Josiah_GP_DMS/htmls/89F_mut_effect.html for a zoomable interactive map of escape mutations across the entirety of GPC. **C** Validation pseudovirus neutralization assays for the indicated GPC mutants against antibody 8.9F. Error bars indicate the standard error for two technical replicates. **D** Correlation of escape scores measured by deep mutational scanning and the IC_50_ measured by pseudovirus neutralization assays. The dashed line represents the highest antibody concentration tested, and so IC_50_s for points on the dashed line are lower bounds. Points are colored as in **C**. The Pearson correlation(r) is indicated.

We first mapped escape mutations from antibody 8.9F, a well-characterized anti-Lassa virus antibody that binds to the apex of GPC.^31^ Escape from 8.9F was concentrated at a small number of sites in GPC that largely fall in the structurally defined epitope (Figure 3B and interactive escape map at https://dms-vep.org/LASV_Josiah_GP_DMS/htmls/89F_mut_effect.html). Some but not all of these mutations are well tolerated for GPC-mediated cell entry (Figure 3B logo plot inset), suggesting the potential for escape from this antibody with little cost to GPC function. The mutation escape values measured by deep mutational scanning were highly correlated with changes in IC_50_ measured in standard pseudovirus neutralization assays (Figure 3C,D), validating the accuracy of the deep mutational scanning escape maps.

We generated escape maps for a total of six neutralizing monoclonal antibodies that were previously isolated from Lassa virus convalescent individuals (Figure 4A and links to interactive escape maps at the beginning of “Methods”). One immediately obvious observation is that all antibodies are escaped by at least some mutations that have little negative impact on GPC-mediated cell entry (see the abundance of darkly colored letters in the logo plots in Figure 4A). This observation suggests that GPC has substantial evolutionary capacity to escape each of these antibodies.

**Figure 4.**
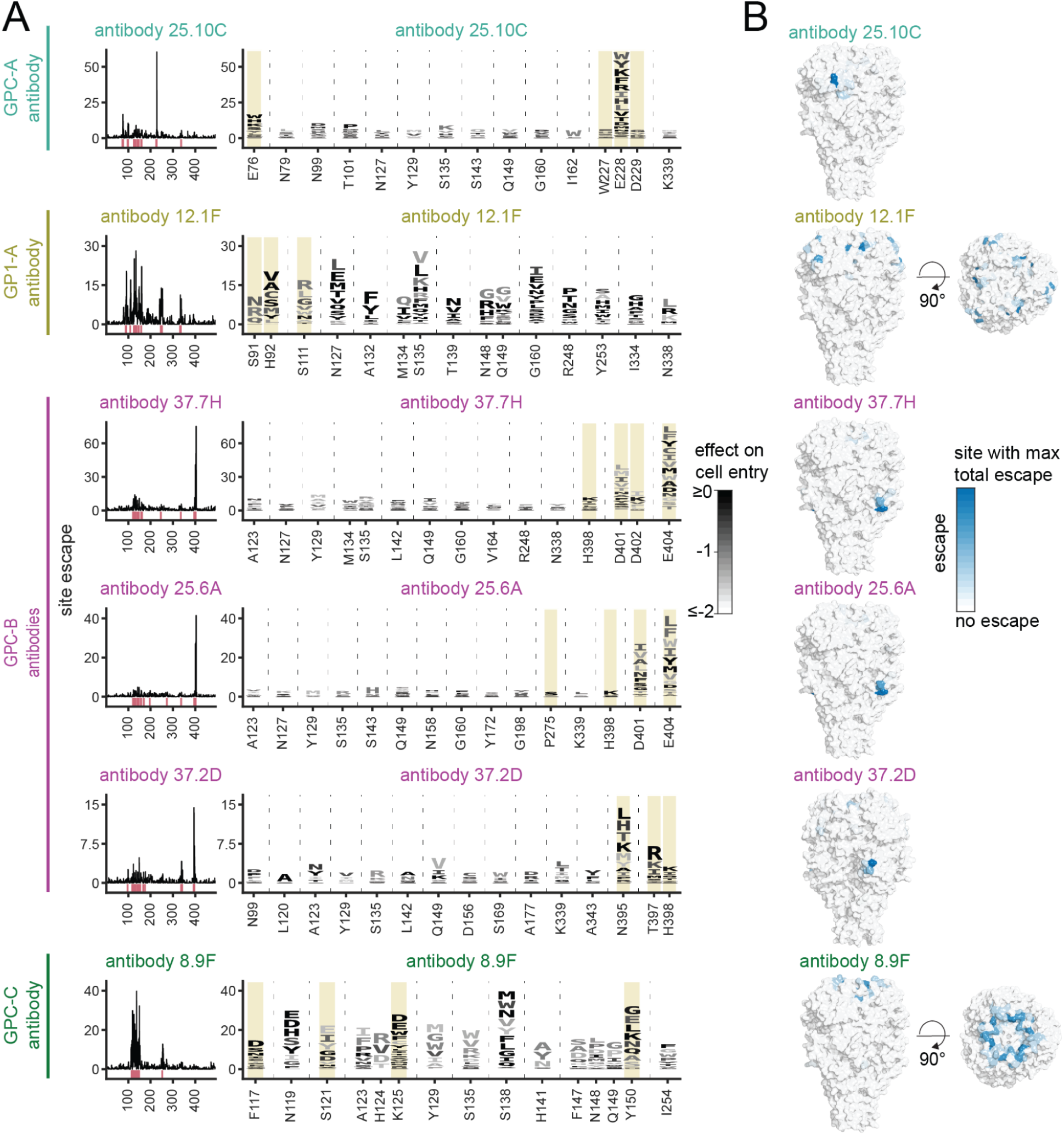
Complete escape maps for six human monoclonal antibodies. **A** Escape maps for each antibody. Line plots show site summed effects of all escape mutations at a site. The top 15 escape sites for each antibody are highlighted pink below the line plot and shown in logo plots where the height of each letter indicates escape caused by that mutation. Letters are colored by mutational effects on cell entry in the absence of antibody, with mutations that impair entry shown in lighter gray. Sites that contact the antibody (within 4 Å) are highlighted in yellow. The antibody escape maps are grouped by the epitope classification in Robinson et al.^28^ **B** Surface representation of Lassa GPC (PDB: 7PUY) colored by summed site escape for each antibody. Blue indicates the site with the most escape from that antibody, and white indicates sites with no escape. See beginning of “Methods” for links to more detailed interactive versions of each escape map.

The escape maps provide high-resolution views that are more informative than the canonical epitope-binning^28^ of GPC antibodies (Figure 4). For instance, antibodies 37.7H and 37.2D both target what has been defined as the GPC-B epitope,^28^ but are escaped by different sets of mutations (e.g., 37.7H is escaped by mutations at site E404 but not site N395, while 37.2D is escaped at site N395 but not site E404). Conversely, antibodies 12.1F and 8.9F recognize what are canonically defined as different epitopes, but are both escaped by mutations at sites S135, N148, and Q149.

Most of the escape mutations occur at sites that fall in the structural footprints of the antibodies as determined by prior high-resolution crystal and cryo-EM structures (Figure 5).^31,48,49,51^ However, only a fraction of the sites in the footprints actually have strong escape mutations (Figures 5 and S7A). In some cases, mutations in the binding footprints are too deleterious to yield GPCs that can facilitate cell entry, and therefore cannot be measured in our assays—for instance, sites L258 and W264 contact antibodies 8.9F and 37.2D, respectively, but are mutationally intolerant (Figure S7A). In other cases, mutations in binding footprints are well tolerated but simply do not affect neutralization—for instance, sites K161 and E100 contact antibodies 12.1F and 25.10C, respectively, and are mutationally tolerant but do not affect neutralization (Figure S7A). The existence of mutationally tolerant sites that contact the antibody but where mutations do not affect neutralization is consistent with prior findings that a small subset of contact residues are primarily responsible for antibody binding energetics and escape.^76,77^

**Figure 5.**
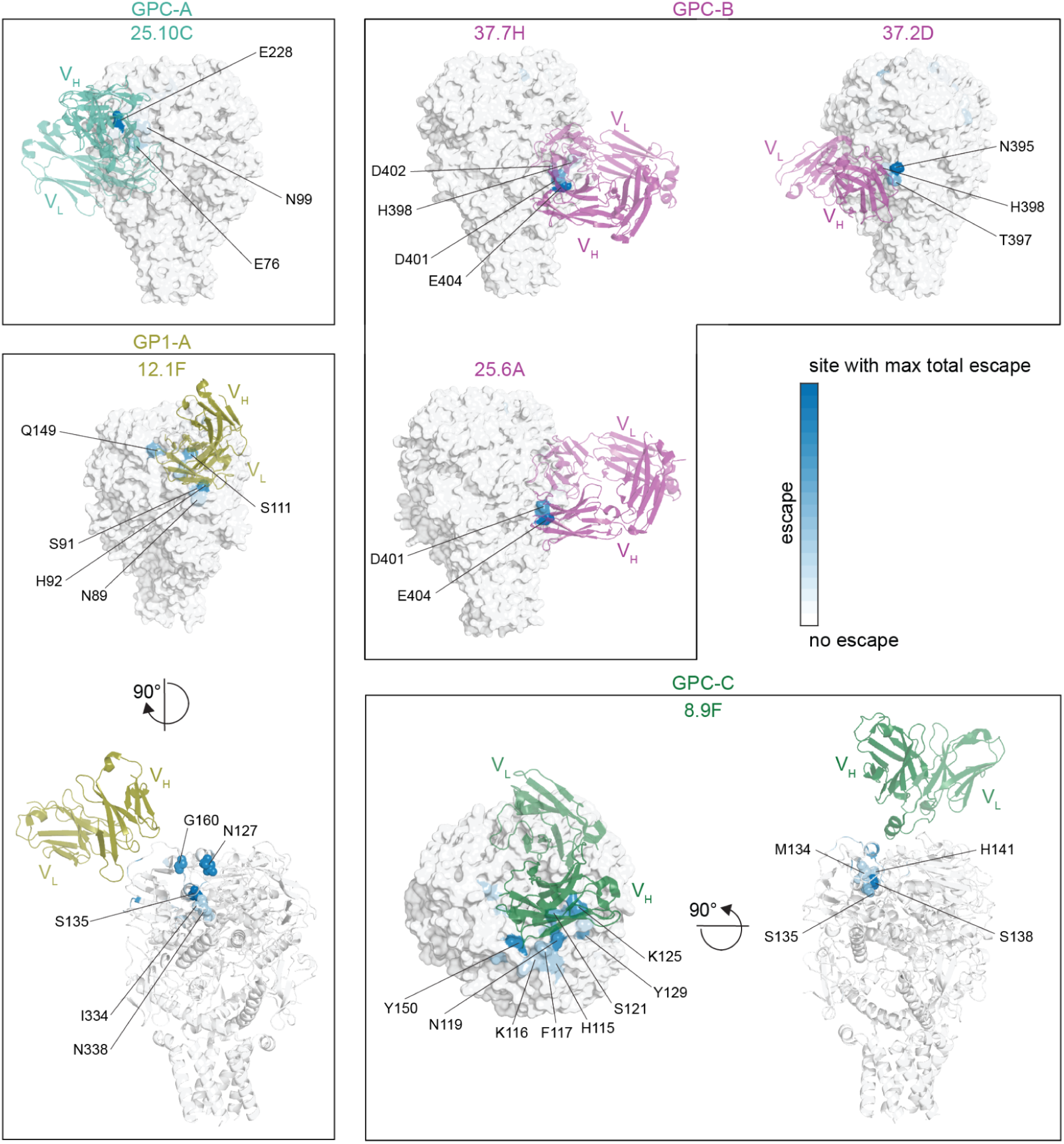
Escape mutations are usually in or near the antibody structural footprint. Surface representation of Fab-bound GPC colored by site escape as measured in deep mutational scanning, with the Fab shown in a colored cartoon representation. Because GPC is a homo-trimer, escape is colored only on sites in the monomer that is closest to the antibody shown for the structure. Blue indicates the GPC site with the most escape from that antibody, and white indicates sites with no escape. The Fab bound antibody structures shown here come from prior crystal and cryo-EM structures.^31,48,49,51^ The antibodies are grouped by epitope as described in Robinson et al.^28^

For some antibodies (e.g., 12.1F and 8.9F), there are strong sites of escape somewhat more distal from the structural binding footprint (Figure 5). In some cases these distal escape sites affect N-linked glycans required for antibody binding (Figure S7B,C). For instance, antibody 12.1F relies on five N-linked glycans (N89, N109, N119, N167, and N224) for binding to GPC, and mutations to the glycosylation motif sites tend to escape 12.1F (Figure S7C).^31,78^ Similarly, antibody 8.9F relies on the N-linked glycan N119 (which is also important for GPC attachment to *ɑ*-DG) and mutations to this glycosylation site escape 8.9F (Figure S7C).^31^ However, glycans do not explain all distal escape sites: for instance, antibody 12.1F is escaped by sites S135, I334, and N338, which cluster together buried below the surface of the GPC bound by the antibody (Figure S7D). Likewise, antibody 8.9F is strongly escaped at sites S138 and H141, which also cluster below the surface of the GPC bound by the antibody.

Three of the antibodies for which we mapped escape (12.1F, 37.2D, and 8.9F) are being developed as an antibody cocktail called Arevirumab-3,^29^ which protects from lethal Lassa virus infection in animal models.^29–32^ To provide insight into how GPC mutations could affect neutralization by the antibodies in this cocktail, we analyzed the escape maps for mutations that reduced neutralization by multiple antibodies in the cocktail (Figure S8). Although the three antibodies have been structurally classified as targeting distinct epitopes,^31^ we identified sites where mutations affect neutralization by multiple antibodies that are components of the cocktail (Figure S8). In particular, antibodies 12.1F and 8.9F are escaped by similar mutations, which emphasizes the importance of antibody 37.2D for making the cocktail robust against viral escape (Figure S8).

### Antibody-escape mutations are present in natural Lassa virus strains

An important factor in designing therapeutic/prophylactic antibodies is assessing how broadly they are expected to neutralize different viral lineages across the current known diversity of Lassa virus. To determine if naturally occurring Lassa virus strains contain GPC mutations that escape the six antibodies we mapped, we searched all available Lassa GPC sequences for mutations that cause escape in our deep mutational scanning data (Figure 6A). For all antibodies, there were strains that naturally contained GPC mutations that escaped neutralization in our deep mutational scanning of the Josiah strain GPC (Figure 6A).

**Figure 6.**
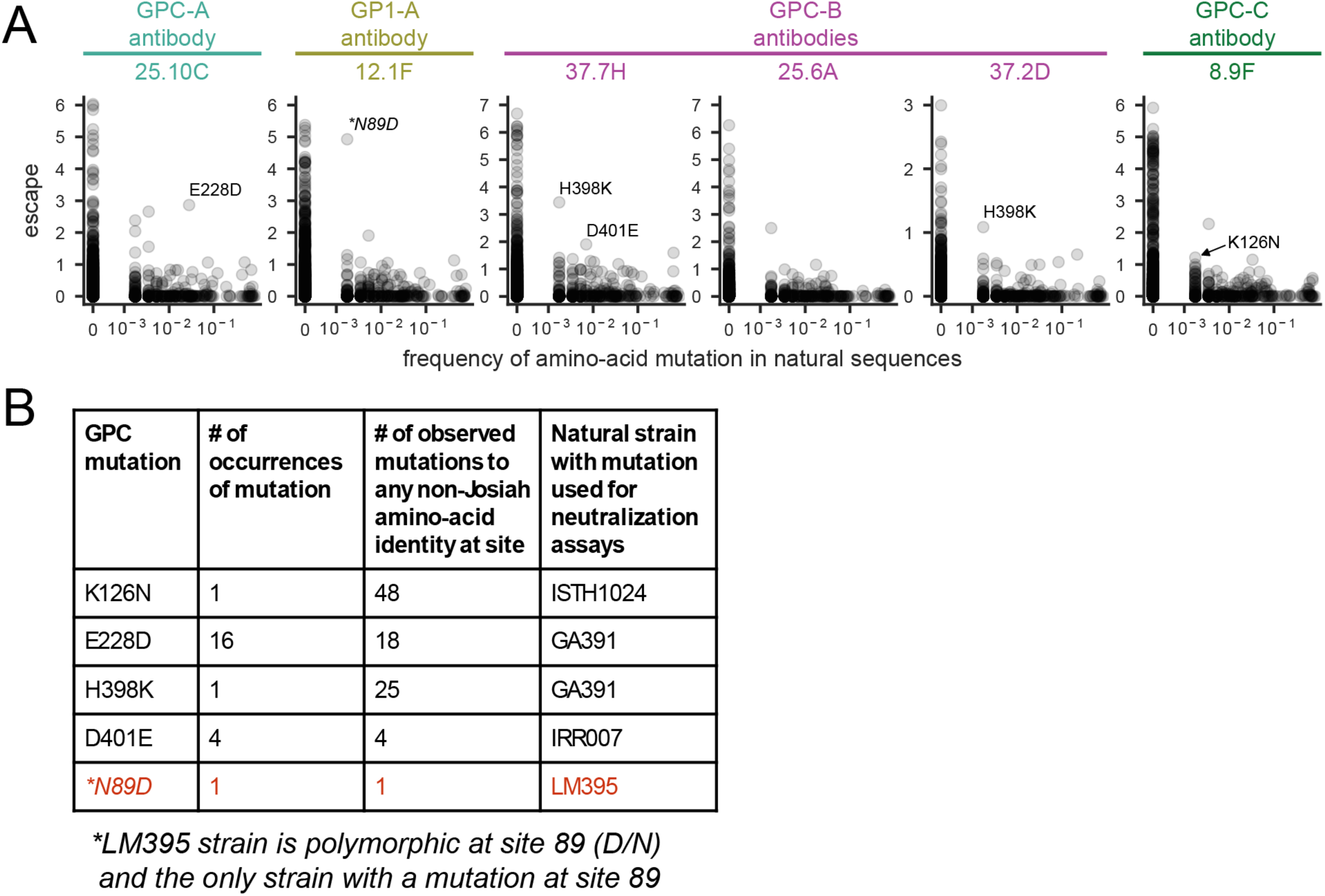
Some antibody escape mutations mapped by deep mutational scanning are found in natural Lassa virus strains. **A** Frequency of mutations that escape antibody neutralization in natural Lassa GPC sequences. The escape for each mutation as measured in deep mutational scanning is plotted versus the mutation’s frequency in all 572 high-quality Lassa GPC sequences. Mutations chosen for validation with pseudovirus neutralization assays are labeled. **B** Summary of escape mutations and representative strain containing each mutation that were chosen for validation experiments in Figure 7. For each mutation, the following are indicated: number of strains containing that mutation, number of strains containing any non-Josiah amino-acid identity at that site, and the natural strain whose GPC contains the mutation that was chosen for testing. N89D is marked with an asterisk because the LM395 is the only strain with a mutation at that site, and the deep sequencing data show site 89 is polymorphic between D and N (Figure S9B).^17^ All mutations shown in this figure are defined relative to the parental amino acid at that site in the Josiah strain GPC.

To validate the increased neutralization resistance of natural strains carrying escape mutations identified in our deep mutational scanning maps, we chose GPCs from four virus strains that contained one or more of these escape mutations (Figures 6B and S9A). (Note that the N89D mutation that escapes antibody 12.1F is a high-frequency polymorphism rather than a fixed mutation in the LM395 strain^17^; Figure S9B). We created pseudoviruses expressing GPC from each strain and performed neutralization assays with each pseudovirus as well as the

Josiah strain containing just the point mutant of interest (Figure 7). Consistent with the deep mutational scanning, each natural GPC variant was more resistant to neutralization by the antibody that it was predicted to escape (Figure 7). This finding confirms that deep mutational scanning of the GPC from the Josiah strain can identify other natural Lassa strains with mutations that escape particular antibodies.

**Figure 7.**
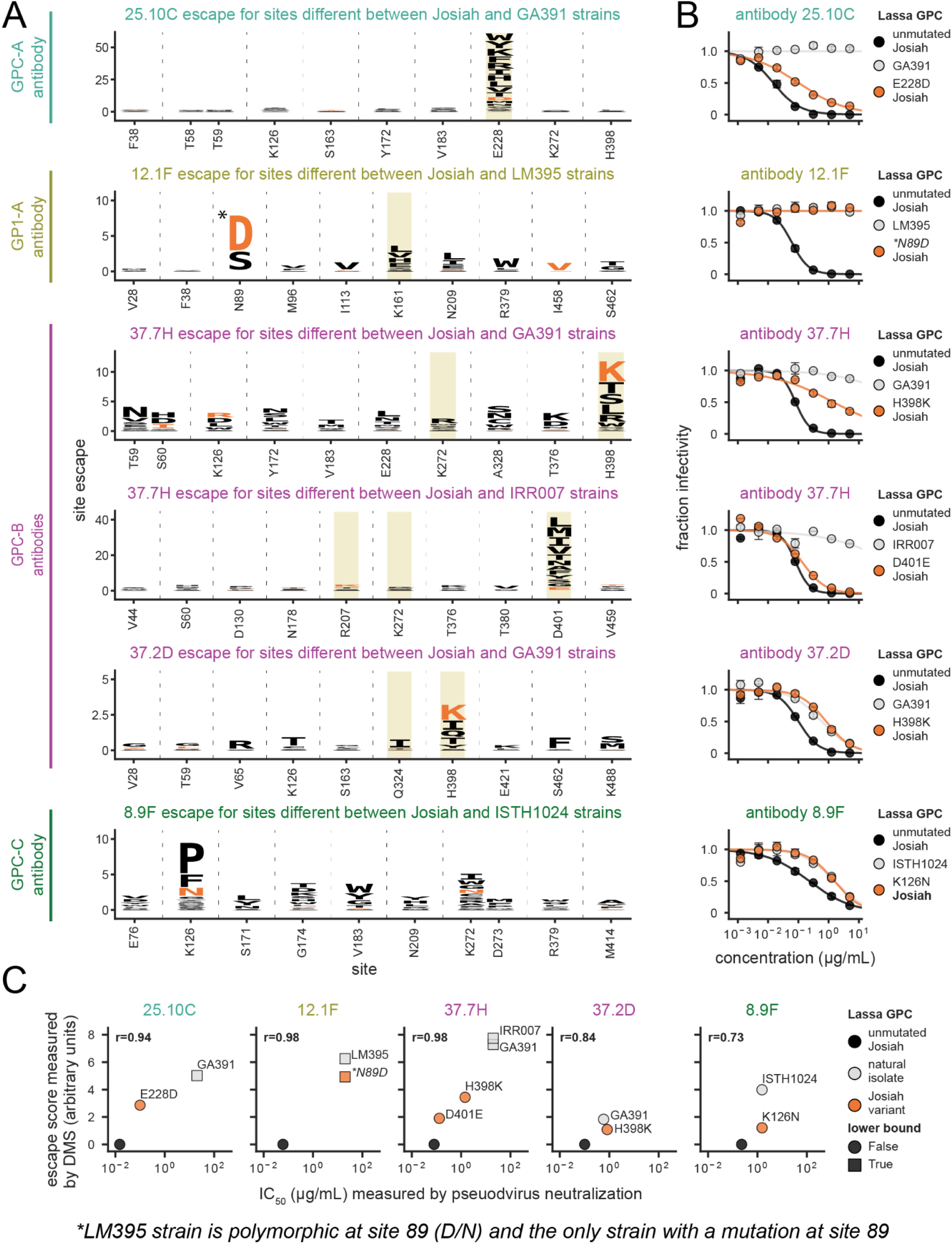
Validation that antibody escape mutations are found in natural GPC variants. **A** Deep mutational scanning antibody escape maps for top 10 escape sites that differ between the Josiah strain GPC and the indicated strain’s GPC for each antibody. The height of the letter corresponds to the strength of escape, and amino acids representing mutations that are found in the indicated strain’s GPC are colored orange (e.g., the third plot from top indicates H398K is found in the GA391 strain). Sites that contact the antibody in structures of the GPC-Fab complex (within 4 Å) are highlighted yellow. The antibody escape maps are grouped by the antibody epitope classification of Robinson et al.^28^ **B** Validation pseudovirus neutralization assays for the indicated Josiah GPC mutants and natural strain GPCs. Unmutated Josiah GPC is colored black, single mutant Josiah GPC is colored orange, and natural strain GPC is colored gray. Error bars indicate the standard error for two technical replicates. **C** Correlation of escape predicted for natural strain GPCs by summing of the effects measured in deep mutational scanning for all mutations in that GPC versus the actual IC_50_ measured by pseudovirus neutralization assays. Squares indicate that the antibody did not neutralize at the highest concentration tested, and so the reported IC_50_ is a lower bound. *N89D is marked because the LM395 strain with the N89D mutation is polymorphic at site 89 (Figure S9B)^17^ and is the only strain with a mutation at site 89.

For several of the naturally occurring GPCs with reduced neutralization, the extent of antibody escape is very similar to that of the corresponding single escape mutant of the Josiah strain GPC we tested (e.g., N89D and the LM395 strain, H398K and the GA391 strain, and K126N and the ISTH1024 strain in Figure 7B). Therefore, for these particular natural GPCs and antibodies, the single mutation identified in deep mutational scanning largely dictates the antibody escape. However, several other natural GPCs showed antibody escape that exceeded that of the corresponding single mutant of the Josiah GPC. For example, strain GA391 completely escaped neutralization from antibody 25.10C, but the strongest escape mutant in the GA391 strain (E228D) only caused partial escape from this antibody in the Josiah strain GPC—suggesting that other mutations in the GA391 GPC also contribute to escape from antibody 25.10C (Figure S10). Indeed, the escape by a given natural strain is well predicted by the sum of the effects of its individual mutations relative to the Josiah strain GPC as measured in the deep mutational scanning (Figure 7C). Therefore, it appears we can predict the escape of natural strains simply by summing the effects of their constituent GPC mutations as measured in the deep mutational scanning.

## Discussion

We have used pseudovirus deep mutational scanning to measure how nearly all mutations to Lassa virus GPC affect cell entry and antibody neutralization. These measurements can accelerate countermeasures by comprehensively assessing how GPC’s genetic diversity impacts neutralization by antibodies, as well as identifying sites in GPC that are likely to be tolerant and intolerant of future evolutionary change. In addition, because our experimental system uses pseudoviruses that can undergo only one round of infection, it enables high-throughput study of GPC mutations at biosafety-level-2 even though authentic Lassa virus is a biosafety-level-4 pathogen.^46^

Our maps of how mutations affect cell entry define the functional constraint throughout GPC. Previous GPC mutational studies^79,80^ have examined just small subsets of mutations, whereas our study maps the impact of nearly all 9,820 GPC amino-acid mutations. These maps reveal that the SSP and GP2 domains (which primarily comprise the GPC stalk) are more functionally constrained than GP1 (which comprises most of the head of the GPC ectodomain). However, within the the head of GPC there are regions of high constraint, including, the core binding pocket for the primary host receptor, *ɑ*-DG. However, the putative binding region^78,79^ to the secondary LAMP1 receptor is more mutationally tolerant, consistent with LAMP1 not being essential for infection^79,90–92^ and different Lassa lineages have varying dependencies on LAMP1 for cell entry.^84^

One potential use of the maps of functional constraint is the design of GPC vaccine immunogens. The development of Lassa virus vaccines have been hindered by the metastable nature of GPC.^48,50,85–87^ Maps of functional constraint throughout GPC can inform the engineering of stable immunogens in several ways. First, these measurements provide insight into where mutations can be introduced without disrupting GPC folding. Second, functionally constrained regions identified in our maps provide attractive targets for structure-guided vaccines designed to elicit antibody responses that will broadly recognize both current and potential future antigenic variants of Lassa virus.

We also directly mapped how GPC mutations affected neutralization by a panel of six human monoclonal antibodies, three of which are being developed as an antibody cocktail called Arevirumab-3.^29^ We found that all six antibodies are partially or completely escaped by mutations that are present in at least some known natural Lassa virus strains. This highlights how development of antibodies for use against Lassa virus should explicitly consider the potential for viral escape in order to avoid the problems that have plagued antibody-based countermeasures against other viruses like SARS-CoV-2.^38–41^ By prospectively mapping potential escape mutations, our work provides a road map to guide the development of anti-Lassa antibodies that will be more robust to both current GPC variation and possible future evolution.

## Limitations of study

Our study uses GPC-pseudotyped lentiviruses to measure the effect of GPC mutations on single-cycle entry into 293T cells and neutralization of entry by antibodies. This approach has the advantage of eliminating the biosafety concerns associated with experiments involving replication competent Lassa virus, but comes with the caveat that pseudovirus cell entry is only a partial proxy for the function of GPC in authentic viral infections.^54,88–90^ As a result, mutations that are favorable or deleterious for cell entry in our experimental system might not always have the same effects on the fitness of actual Lassa virus, although pseudovirus deep mutational scanning measurements have proven informative for understanding the fitness of SARS-CoV-2 variants.^44^ We do note, however, that there is extensive literature indicating that pseudovirus neutralization titers usually correlate well with those measured using authentic virus.^45^

We measured the effects of mutations in the GPC from just a single Lassa virus strain (the Josiah strain). Although we showed that our antibody escape maps were fairly predictive of the antigenic effects of mutations in other diverged GPCs, mutations can have strain-dependent effects due to epistasis among multiple mutations.^64,91^ Therefore, caution should always be used in extrapolating the effects of mutations to different strains.

As with all experiments, our deep mutational scanning measurements involve some noise. The interactive plots described in the first section of the Methods provide several metrics (such as reproducibility of specific measurements across libraries / replicates, and the number of library variants in which a specific mutation is seen) that can be used to assess the accuracy of measurements for specific mutations, and we encourage readers making detailed use of our data to familiarize themselves with these metrics and examine them for mutations of interest.

## Acknowledgements

We thank Dr. Sean Whelan for providing the 293TΔDAG1 cell line. We thank Brendan Larsen and Bernadeta Dadonaite for helpful comments on the manuscript. This work was supported in part by the NIH/NIAID under grant R01AI141707 to J.D.B. K.H.D.C. was supported by NIH/NIAID grant F30AI149928. K.G.A was supported by NIH grants U19AI135995, U01AI151812, and UL1TR002550. J.D.B. and F.A.M. are investigators of the Howard Hughes Medical Institute. This research was also supported by the Genomics & Bioinformatics Shared Resource, RRID:SCR_022606, of the Fred Hutch/University of Washington Cancer Consortium (P30 CA015704), by the Flow Cytometry Shared Resource, RRID:SCR_022613, of the Fred Hutch/University of Washington/Seattle Children’s Cancer Consortium (P30 CA015704), and by Fred Hutch Scientific Computing, NIH grants S10-OD-020069 and S10-OD-028685.

## Methods

### Data availability and interactive plots of results

All code used for analysis in this study is publicly available on GitHub (https://github.com/dms-vep/LASV_Josiah_GP_DMS). An easy to view HTML summary of the analyses with interactive plots of the results is provided on GitHub Pages (https://dms-vep.github.io/LASV_Josiah_GP_DMS/).

We recommend using the interactive plot at https://dms-vep.org/LASV_Josiah_GP_DMS/htmls/293T_entry_func_effects.html to visualize the effects of mutations on cell entry. The visualization includes a site level zoom bar that allows selection of specific GPC sites, a site level line plot, and a mutation specific heatmap. Both the line plot and heatmap can be moused over to display additional information for a specific site or mutation. Importantly, mousing over the heatmap shows the measured value for each experimental replicate for each library—measurements that have consistent values among replicates and libraries are more likely to be accurate. Below the heatmap are several options to change the metrics displayed. The site level line plot can be changed to show either the site sum, mean, maximum, or minimum of mutation effects at that site. The measurements can also be floored at zero to show all negative values as zero. There are also four sliders for the following statistics that filter the data displayed in the line plot and heatmap:

● *minimum times_seen*: number of unique variants (averaged across both libraries) that have a specific mutation. The more variants in which a mutation is measured, the more accurate the measurement is likely to be.
● *minimum n_selections*: number of unique experiments where a given mutation was measured.
● *maximum effect_std*: maximum standard deviation of a mutation measurement across the selection experiments.
● *minimum max of effect at site*: minimum mutation measurement measured at a site. This option can be used to zoom into sites with the most favorable mutation effects..

The data on mutational effects on cell entry displayed in the interactive visualization and filtered using the default values shown in that visualization are available at https://github.com/dms-vep/LASV_Josiah_GP_DMS/blob/ main/results/func_effects/averages/293T_entry_func_effects.csv in CSV format.

We recommend the following interactive visualizations of the effects of mutations on antibody escape for the easy exploration of data.

● 37.7H: https://dms-vep.org/LASV_Josiah_GP_DMS/htmls/377H_mut_effect.html
● 8.9F: https://dms-vep.org/LASV_Josiah_GP_DMS/htmls/89F_mut_effect.html
● 25.10C: https://dms-vep.org/LASV_Josiah_GP_DMS/htmls/2510C_mut_effect.html
● 12.1F: https://dms-vep.org/LASV_Josiah_GP_DMS/htmls/121F_mut_effect.html
● 25.6A: https://dms-vep.org/LASV_Josiah_GP_DMS/htmls/256A_mut_effect.html
● 37.2D: https://dms-vep.org/LASV_Josiah_GP_DMS/htmls/372D_mut_effect.html

Similar to the visualizations for cell entry, the antibody escape visualizations include a site level zoom bar that allows selection of specific GPC sites, a site level line plot, and a mutation specific heatmap. Both the line plot and heatmap can be moused over to display additional information for a specific site or mutation, including most crucially the measured value in different replicates / libraries. Note the heatmaps distinguish between mutations that are not present in the libraries (light gray) versus mutations that are present but are so deleterious for cell entry that their effect on antibody escape cannot be measured (dark gray boxes). Below the heatmap are several options to change the metrics displayed. The site level line plot can be changed to show either the site sum, mean, maximum, or minimum of mutation effects. The measurements can also be floored at zero to show all negative values as zero. There are also five sliders for the following statistics that filter the data displayed in the line plot and heatmap:

● *minimum times_seen*: number of unique variants (averaged across both libraries) that have a specific mutation. Larger values generally indicate a more accurate measurement.
● *minimum functional effect*: minimum effect on cell entry for a given mutation to be displayed for antibody escape. If a mutation has an effect on cell entry less than this minimum, then it is displayed as a dark gray box. The rationale for filtering mutations that have deleterious effects on cell entry is that a mutation must still result in a GPC that can facilitate entry into cells in order to accurately measure its effect on antibody neutralization. Mutations that have effects on cell entry less than the minimum cutoff are denoted in the downloadable CSV file (see link below) by the *poor_cell_entry* column.
● *minimum n_models*: number of unique experiments where a given mutation was measured.
● *maximum escape_std*: maximum standard deviation of a mutation measurement across the selection experiments.
● *minimum max of escape at site*: minimum mutation measurement measured at a site. This slider is useful for zooming in on the most important sites of escape.

The data on antibody escape filtered using the default values in the interactive visualizations are at https://github.com/dms-vep/LASV_Josiah_GP_DMS/tree/main/results/filtered_antibody_escape_CSVs in CSV format.

Note that numerous other data files and interactive plots are also available via the main GitHub Pages link (https://dms-vep.org/LASV_Josiah_GP_DMS/).

### Plasmid maps and primer sequences

All plasmid maps for plasmids used in this study are found in a subdirectory (https://github.com/dms-vep/LASV_Josiah_GP_DMS/tree/main/non-pipeline_analyses/plasmid_maps) in the GitHub repository. Similarly, all primer sequences for primers used in this study are found (https://github.com/dms-vep/LASV_Josiah_GP_DMS/blob/main/non-pipeline_analyses/primers/primers.csv) in the GitHub repository.

### Cell lines

293T cells were from ATCC (CRL-3216) and 293T-rtTA expressing cells were the same cell clone previously described in Dadonaite et al.^42^ 293TΔDAG1^27^ cells were a kind gift from Dr. Sean Whelan. To produce 293TΔDAG1 expressing either human or mastomys *DAG1*, we first generated VSV-G pseudotyped lentivirus carrying either the human or mastomys *DAG1* gene. To produce these viruses, we plated 1 million 293T cells per well of a 6-well tissue culture dish. Approximately, 24 hours after plating the cells, we transfected 1 µg of lentivirus backbone containing either human (3699_pHAGE2_EF1a_human_DAG1_IRES_mCherry.gb) or mastomys (3700_pHAGE2_EF1a_mastomys_DAG1_IRES_mCherry.gb) DAG1, 0.25 µg of each lentivirus helper plasmid (Gag-Pol, Tat, and Rev that correspond to the following plasmid maps: 26_HDM_Hgpm2.gb, 27_HDM_tat1b.gb, and 28_pRC_CMV_Rev1b.gb), and 0.25 µg of VSV-G expression plasmid (29_HDM_VSV_G.gb) per well using BioT (Bioland Scientific, Cat. No. B01-02) following manufacturer’s protocol and recommendations for reagent ratios. Note this lentiviral backbone is self-inactivating because of a U3 deletion in the 3’ LTR sequence and that the full length mastomys *DAG1* is from *Mastomys coucha* because this is the only fully annotated *DAG1* sequence available for the *Mastomys* genus. It has previously been shown that *ɑ*-DG is 100% identical at the amino-acid level between *Mastomys natalensis* and *Mastomys coucha*^95^ and only *ɑ*-DG affects Lassa virus GPC attachment (not β-DG).^96^ The supernatant was collected 48 hours after transfection and passed through a 0.45 µM SFCA syringe filter (Corning, Cat. No. 431220). The rescued virus was titrated on 293T cells using protocols described in Crawford et al.^73^ After titrating the lentivirus, we infected 293TΔDAG1^27^ cells at an MOI of 0.1. Approximately, 48 hours post infection single cell clones were sorted into a 96-well plate using BD FACSAria II cell sorter based on mCherry expression from the lentiviral backbone. Single clones were expanded and tested for infectability by GPC-pseudovirus and *ɑ*-DG surface expression as described below. The specific clones chosen and used are shown in Figure S3.

All cell lines were grown in D10 media (Dulbecco’s Modified Eagle Medium with 10% heat-inactivated fetal bovine serum, 2 mM l-glutamine, 100 U/mL penicillin, and 100 µg/mL streptomycin). To avoid rtTA activation and mutant GPC expression earlier than intended, 293T-rtTA cells were grown in D10 made with tetracycline-free fetal bovine serum (Gemini Bio, Cat. No. 100-800). For generation of GPC- and VSV-G-pseudotyped variant libraries, D10 media was made with phenol-free DMEM (Corning DMEM With 4.5g/L Glucose, Sodium Pyruvate; Without L-Glutamine, Phenol Red from Fisher, Cat. No. MT17205CV). Note that the VSV-G-pseudotyped variant libraries used for PacBio sequencing described below were produced in D10 media (Dulbecco’s Modified Eagle Medium with 10% heat-inactivated fetal bovine serum, 2 mM l-glutamine, 100 U/mL penicillin, and 100 µg/mL streptomycin).

### Flow cytometry analysis of ɑ-DG surface expression

To measure the cell surface expression of *ɑ*-DG in the various cell lines as shown in Figure S3B, we plated approximately 5 million 293T, 293TΔDAG1, 293TΔDAG1+humanDAG1, and 293TΔDAG1+mastomysDAG1 cells in 10 cm tissue culture plates. Approximately, 24 hours later, we removed the cells from the dish using Gibco Cell Dissociation Buffer, enzyme-free, Hanks’ Balanced Salt Solution (ThermoFisher, Cat. No. 13150016). The cells were washed with FACS buffer (PBS + 1% bovine serum albumin (BSA)) and 1 mL aliquots of cells were incubated with 1 µg of sheep anti-human dystroglycan antibody (R&D Systems, Cat. No. AF6868) for 30 minutes on ice. Due to the conservation of human and mastomys *ɑ*-DG as shown in Figure S2, an anti-human dystroglycan antibody was used to stain for expression in both 293TΔDAG1+humanDAG1 and 293TΔDAG1+mastomysDAG1 cell lines. For a no-staining control, cells were incubated on ice without any antibody. After the incubation, the cells were washed with FACS buffer two times and resuspended in 1 mL. After washing, the cells were incubated with 5 µL (1:200) donkey anti-sheep IgG APC-conjugated antibody (R&D Systems, Cat. No. F0127) for 30 minutes on ice. After the incubation, the cells were washed with FACS buffer two times and resuspended in 1 mL of FACS buffer. The cells were analyzed with a BD FACSymphony A5 cytometer and the data were plotted using FlowJo software (Version 10, BD Biosciences, Ashland, OR, USA).

### Design of lentiviral vector backbone containing Lassa GPC

An overview of the lentiviral backbone is shown in Figure 1A and plasmid map (2912_pH2rU3_ForInd_LASV_Jos-re-opt_CMV_ZsG_fixK.gb) is provided in the GitHub repository. The backbone has a repaired 3′ LTR allowing for re-rescue after integration into cells,^92^ constitutive expression of zsGreen as a selectable marker for infection, and a TRE3G promoter that inducibly expresses GPC when the reverse tetracycline transactivator (rtTA) in the 293T-rtTA cells is induced by the presence of doxycycline.^42^ The GPC sequence used in the backbone is a codon optimized Josiah strain GPC.^47^ Unlike in Dadonaite et al,^42^ the lentiviral backbone used here does not have the puromycin resistance gene. The reason is that the current study was actually started before Dadonaite et al^42^; for future studies we suggest using the backbone in Dadonaite et al^42^ that contains the puromycin resistance gene rather than the backbone used here.

### Design of primers for Lassa GPC mutagenesis

Our goal was to perform site saturation mutagenesis of all sites in the Josiah strain GPC. To mutagenize every site, we used primers tiled along the entire gene that contained an NNS codon. We used an NNS mutagenesis strategy because of the reduced frequency of stop codons and the more balanced representation of amino acids.^97^ Primer sequences for all mutations were generated using a Python script (https://github.com/jbloomlab/TargetedTilingPrime <u>rs</u>) and ordered as oPools from Integrated DNA Technologies. See https://github.com/dms-vep/LASV_Josiah_GP_D <u>MS/tree/main/non-pipeline_analyses/library_construction</u>for all code related to primer design. Codon tiling primer sequences (https://github.com/dms-vep/LASV_Josiah_GP_DMS/blob/main/non-pipeline_analyses/library_constr <u>uction/design/codon_tiling_primers/20210114_LASVGP_Josiah_reopt_AllPrimers.xlsx</u>) can be found in the GitHub repository.

### Production of Lassa GPC deep mutational scanning plasmid libraries

The general workflow for producing the deep mutational scanning plasmid libraries consisted of mutagenizing the GPC sequence, barcoding the mutagenized GPC sequences, and cloning the mutagenized/barcoded GPC sequences into the lentiviral backbone. We created two independent biological library replicates (library A and B) by performing two independent plasmid preps of the following steps. As a result, each library has a unique set of mutations and barcodes.

To mutagenize the GPC sequence, a codon optimized GPC sequence was first amplified from a plasmid containing the codon optimized sequence in a lentiviral backbone (2912_pH2rU3_ForInd_LASV_Jos-re-opt_CMV_ZsG_fixK.gb). The PCR was performed using 1.5 µL of 10mM forward primer (VEP_amp_for), 1.5 µL of 10 mM reverse primer (3’rev_lib_LinJoin_KHDC), 10 ng of GPC gene template, 25 µL of KOD polymerase (KOD Hot Start Master Mix, Sigma-Aldrich, Cat. No. 71842), and water for a final volume of 50 µL. PCR cycling conditions were:

1. 95°C for 2 min
2. 95°C for 20 s
3. 70°C for 1 s
4. 58°C for 10 s, cooling at 0.5°C per 1 s
5. 70°C for 40 s (return to step 2 for another 19x cycles)
6. Hold at 4°C

The amplified, linearized GPC sequence was gel purified using NucleoSpin Gel and PCR Clean-up kit (Takara, Cat. No. 740609.5) and then purified using Ampure XP beads (Beckman Coulter, Cat. No. A63881) at 1:1 sample to bead ratio.

Next, the purified GPC template was used in a mutagenesis PCR using a similar protocol described previously in Bloom.^53^ Forward and reverse pools of NNS codon tiling primers (https://github.com/dms-vep/LASV_J <u>osiah_GP_DMS/blob/main/non-pipeline_analyses/library_construction/design/codon_tiling_primers/20210114_LASV</u> <u>GP_Josiah_reopt_AllPrimers.xlsx</u>) for generating mutations were produced as described above. The forward and reverse primer pools were combined with the reverse linearizing primer (3’rev_lib_LinJoin_KHDC) and forward linearizing primer (VEP_amp_for), respectively in separate reactions. Each PCR reaction was performed using 8 µL H2O, 1.5 µL DMSO, 4 µL 3 ng/µL linearized GPC template, 1.5 µL 5 µM forward or reverse primer pool, 1.5 µL reverse (3’rev_lib_LinJoin_KHDC) or forward (VEP_amp_for) linearizing primer, and 15 µL 2× KOD Hot Start Master Mix. PCR cycling conditions were:

1. 95°C for 2 min
2. 95°C for 20 s
3. 70°C for 1 s
4. 54°C for 20 s, cooling at 0.5°C per 1 s
5. 70°C for 50 s (return to step 2 for another 9x cycles)
6. Hold at 4°C

Next, the forward and reverse mutagenized PCR products were joined. The joining PCR was performed using 4 µL H2O, 4µL of 1:4 diluted forward mutagenic reaction, 4 µL 1:4 diluted reverse mutagenic reaction, 1.5 µL 5 µM (VEP_Amp_For), 1.5 µL 5 µM (3’rev_lib_LinJoin_KHDC), 15 µL 2x KOD Hot Start Master Mix. PCR cycling conditions were:

1. 95°C for 2 min
2. 95°C for 20 s
3. 70°C for 1 s
4. 54°C for 20 s, cooling at 0.5°C per 1 s
5. 70°C for 50 s (return to step 2 for another 19x cycles)
6. Hold at 4°C

To barcode the mutagenized GPC sequences, the PCR products from the joining PCR reaction were first gel and Ampure XP purified. The purified GPC sequences were then barcoded with random 16-nucleotide barcodes downstream of the GPC stop codon.^42^ Barcodes with 16 N nucleotides were chosen because this allows for 4^16^ possible barcodes, which is much greater than the size of our deep mutational scanning plasmid libraries. The large diversity of barcodes reduces the chances of a barcode being duplicated between two different variants. The barcoding PCR was performed using 9 µL H2O, 30 ng of joining PCR product, 1 µL 10uM (5’_BC), 1 µL 10uM (ForInd_AddBC_2), 15 µL 2x KOD Hot Start Master Mix. Cycling:

1. 95°C for 2 min
2. 95°C for 20 s
3. 70°C for 1 s
4. 55.5°C for 20 s, cooling at 0.5°C per 1 s
5. 70°C for 55 s (return to step 2 for another 9x cycles)
6. Hold at 4°C

To clone the mutagenized and barcoded GPC sequences into the lentiviral backbone, the PCR products from the barcoding PCR reaction were first gel and Ampure XP purified. The mutagenized GPCs were then cloned into the lentiviral backbone by first digesting the backbone plasmid (2871_pH2rU3_ForInd_mCherry_CMV_ZsG_NoBC_cloningvector.gb) at the Mlul and Xbal sites. The digested backbone was gel and Ampure XP purified. We then assembled the digested backbone with the mutagenized GPC inserts with NEBuilder HiFi DNA Assembly kit (NEB, Cat. No. E2621) at a 1:2 vector to insert ratio for a 1 hr reaction. The HiFi products were gel and Ampure XP purified. We used 1 µL of purified HiFi product to transform 20 µL of

10-beta Electrocompetent *E. coli* cells (NEB, Cat. No. C3020K). We performed 4 electroporation reactions for a final count of ∼300,000 CFUs per library and plated transformed cells on LB+ampicillin plates. To reduce barcode duplication as explained in the next section, we aimed to create plasmid libraries from a much larger number of CFUs than the final number of variants in our virus libraries.^42^ The plates were incubated overnight at 37°C and scraped plates into liquid LB. Plasmid stocks were prepared using ten separate 1.5 mL minipreps (QIAprep Spin Miniprep Kit, Cat. No. 27106X4).

### Production of cells that contain integrated Lassa GPC variants

The general workflow for producing cells integrated with single barcoded GPC variants involved producing VSV-G pseudotyped lentiviruses that contain GPC variants in their genomes, infecting rtTA-expressing 293T cells with VSV-G pseudotyped viruses, and selecting for transduced cells (Figure 1B).

To create the VSV-G pseudotyped viruses, we aimed to produce many more transducing units than the number of colonies scrapped during cloning to reduce any bottlenecks in barcoded variant diversity. We plated 0.5 million 293T cells per well in nine wells of two 6-well tissue culture plates for each library. Approximately 24 hours later, cells were transfected using BioT (Bioland Scientific, Cat. No. B01-02) following manufacturer’s protocol and recommendations for reagent ratios. we transfected 0.25 µg of each lentivirus helper plasmids (Gag-Pol, Tat, and Rev that correspond to the following plasmid maps: 26_HDM_Hgpm2.gb, 27_HDM_tat1b.gb, and 28_pRC_CMV_Rev1b.gb), 0.25 µg VSV-G expression plasmid (29_HDM_VSV_G.gb), and 1 µg lentivirus backbone containing mutagenized, barcoded GPC (described in previous section). Media was changed ∼24 hours post transfection and viruses were harvested ∼48 hours after the media was changed. To harvest the virus, the transfection supernatants were collected and filtered through a 0.45 µM SFCA filter (Corning, Cat. No. 431220). Viruses were stored at −80°C and titrated as described in Crawford et al.^73^

Next, we infected rtTA-expressing 293T cells^42^ with the approximate number of VSV-G-pseudotyped transducing units carrying barcoded GPC variants as the desired number of variants in final libraries, which was ∼50,000 barcoded variants per library. Note that we infected cells with a lower number of variants compared to the possible number of variants present in our plasmid libraries because we wanted to reduce barcode duplication from recombination in the lentivirus genome.^42,98–100^ Cells were infected at an MOI < 0.02 because we wanted only a single GPC variant integrated into each cell. The transduced rtTA-expressing 293T cells were selected for using fluorescence-activated cell sorting by detecting ZsGreen expression from the lentiviral backbone. In addition, when selecting for the transduced cells, the targeted low MOI was confirmed. The sorted cells were expanded and then cell aliquots were frozen in D10 with tetracycline-free FBS (Gemini Bio, Cat. No. 100-800) containing 10% DMSO. Frozen cell aliquots were stored in liquid nitrogen.

### Generation of Lassa GPC pseudotyped and VSV-G pseudotyped lentiviral particles

To produce genotype-phenotype linked Lassa GPC-pseudotyped lentiviruses, we plated 75 million integrated library cells (described in previous section) per 5-layer flask (Corning Falcon 875cm^2^ Rectangular Straight Neck Cell Culture Multi-Flask, Cat. No. 353144) in 150 mL of D10 without phenol red supplemented with 100 ng/mL of doxycycline (which induces GPC expression ahead of pseudovirus production) per library. Approximately 24 hours after plating, the cells were transfected with 50 µg of each lentivirus helper plasmids (Gag-Pol, Tat, and Rev that correspond to the following plasmid maps: 26_HDM_Hgpm2.gb, 27_HDM_tat1b.gb, and 28_pRC_CMV_Rev1b.gb) using BioT (Bioland Scientific, Cat. No. B01-02) following manufacturer’s protocol and recommendations for reagent ratios. The supernatant was collected 48 hours after transfection and passed through a 0.45 µM SFCA Nalgene 500mL Rapid-Flow filter unit (Cat. No. 09-740-44B). Filtered supernatant was concentrated approximately 10-fold by spinning at 4°C 2000 RCF for 30 min using a Pierce Protein Concentrator (ThermoFisher, Cat. No. 88537) and then virus aliquots were stored at −80°C and titrated as described in Crawford et al.^73^ The titers of the concentrated GPC-pseudotyped virus were typically in the range of 1.5×10^5^ to 2×10^5^ transduction units per mL. Concentrating the GPC-pseudotyped variant library virus was essential because of the presence of deleterious mutations that lowered titers.

To produce VSV-G pseudotyped virus to use for PacBio sequencing to link GPC mutations to barcodes, we plated 15 million integrated library cells per 15 cm tissue culture dish in 30 mL of D10 per library. Approximately 24 hours after plating, the cells were transfected with 7.5 µg of each lentivirus helper plasmids (Gag-Pol, Tat, and Rev that correspond to the following plasmid maps: 26_HDM_Hgpm2.gb, 27_HDM_tat1b.gb, and 28_pRC_CMV_Rev1b.gb) and 7.5 µg of VSV-G expression plasmid (29_HDM_VSV_G.gb) using BioT (Bioland Scientific, Cat. No. B01-02) following manufacturer’s protocol and recommendations for reagent ratios. The supernatant was harvested 48 hours after transfection and filtered through a 0.45 µM SFCA filter (Corning, Cat. No. 431220). Virus aliquots were stored at −80°C and titrated as described in Crawford et al.^73^

To produce VSV-G pseudotyped virus to use as controls for selections to determine effects on cell entry, we plated 15 million integrated library cells per 15 cm tissue culture dish in 30 mL of D10 without phenol red supplemented with 100 ng/mL of doxycycline per library. Approximately 24 hours after plating, the cells were transfected with 7.5 µg of each lentivirus helper plasmids (Gag-Pol, Tat, and Rev that correspond to the following plasmid maps: 26_HDM_Hgpm2.gb, 27_HDM_tat1b.gb, and 28_pRC_CMV_Rev1b.gb) and 7.5 µg of VSV-G expression plasmid (29_HDM_VSV_G.gb) using BioT (Bioland Scientific, Cat. No. B01-02) following manufacturer’s protocol and recommendations for reagent ratios. The supernatant was harvested 48 hours after transfection and filtered through a 0.45 µM SFCA filter (Corning, Cat. No. 431220). Virus aliquots were stored at −80°C and titrated as described in Crawford et al.^73^

### Long-read PacBio sequencing of Lassa GPC barcoded variants

To link GPC mutations with barcodes, we used long-read PacBio sequencing to sequence the entire GPC and random 16-nucleotide barcodes. We sequenced variants produced from the integrated cells because these variants represent the final genotype-phenotype linked variants that will be used for selections. Note that the plasmid libraries cannot be sequenced to link variants and barcodes because of the recombination that occurs to make the cells integrated with GPC variants. We first plated 0.5 million 293T cells per well in a 6-well tissue culture dish. Approximately 24 hours later, we infected three wells each with 1 million VSV-G-pseudotyped variant transducing units per library. Note that the number of transducing units is much greater than the expected number of barcoded variants in each library, which allows for high variant coverage and more accurate mutation calling for each variant. To target non-integrated lentivirus genomes, we trypsinized the cells, washed the cells with PBS, and miniprepped each well using QIAprep Spin Miniprep Kit (Cat. No. 27106X4) approximately 12 hours after infecting the cells.^101,102^ Each miniprepped sample was eluted in 50 µL of EB. We use non-integrated lentivirus genomes as sequencing templates for both PacBio and Illumina sequencing (described below) because they are more abundant than the integrated provirus.^103–105^ However, note that when titrating pseudovirus as shown Figure S1A, we determine the number of transducing units.

Next, a two-step PCR strategy was used to amplify the barcoded GPC variants recovered from the infected cells as previously described in Dadonaite et al.^42^ To allow for detection of strand exchange that may occur during PCR amplification, we performed the first round PCR using primers that contain single nucleotide tags. The first round PCR was performed using the volume of miniprep product for 2.5×10^6^ copies of DNA as analyzed by qPCR, 1 µL of 10 µM 5’ nucleotide tagging primer (PacBio_5pri_G or PacBio_5pri_C), 1 µL of 10 µM 3’ nucleotide tagging primer (PacBio_3pri_C or PacBio_3pri_G), 20 µL KOD Hot Start Master Mix, and remaining volume H2O for a final volume of 40 µL. For future deep mutational scanning libraries, please see the protocol described in Dadonaite et al,^42^ which does not use qPCR to determine the amount of miniprep product to add to the first round PCR. Cycling conditions for first round PCR:

1. 95°C for 2 min
2. 95°C for 20 s
3. 70°C for 1 s
4. 60°C for 10 s, cooling at 0.5°C per 1 s
5. 70°C for 51 s (return to step 2 for another 7x cycles)
6. 70°C for 60 s
7. Hold at 4°C

The PCR products were Ampure XP purified with a sample to bead ratio of 1:1.25. The second round PCR was performed using 10.5 µL of first variant tag set round 1 PCR product, 10.5 µL of second variant tag set round 1 PCR product, 2 µL of 10 µM 5’ PacBio round 2 forward primer (PacBio_5pri_RND2), 2 µL of 10 µM 3’ PacBio round 2 reverse primer (PacBio_3pri_RND2), and 25 µL KOD Hot Start Master Mix. Cycling conditions:

1. 95°C for 2 min
2. 95°C for 20 s
3. 70°C for 1 s
4. 60°C for 10 s, cooling at 0.5°C per 1 s
5. 70°C for 51 s (return to step 2 for another 11x cycles)
6. 70°C for 60 s
7. Hold at 4°C

The PCR products were Ampure XP purified with a sample to bead ratio of 1:1. Each library sample was then barcoded for PacBio sequencing using SMRTbell prep kit 3.0, bound to polymerase using Sequel II Binding Kit 3.2, and then sequenced using a PacBio Sequel IIe sequencer with a 20-hour movie collection time.

To analyze the PacBio long-read sequencing data, we used *alignparse* (https://jbloomlab.github.io/alignpars <u>e/</u>)^106^ as previously described in Dadonaite et al^42^ to link GPC mutations and barcodes. In brief, we first checked for strand exchange that occurred during PCR amplification using the single nucleotide tags mentioned above. Less than one percent of PacBio CCSs had mismatched nucleotide tags, indicating a low rate of strand exchange. Any CCS with mismatched nucleotide tags were removed as well as any CCSs with out of frame indels. We then calculated empirical accuracies of the CCSs by determining how often CCSs with the same barcode had the same GPC sequence. The empirical accuracies were ∼0.8 (the fraction of CCSs correctly reporting the actual mutations). The inaccuracies are due to sequencing errors, strand exchange, reverse transcription errors, and actual barcode duplication where different variants have the same barcode. Consensus sequences were then created using a minimum requirement of three CCSs per variant-barcode sequence to create a barcode-variant lookup table (https://github.com/dms-vep/LASV_Josiah_GP_DMS/blob/main/results/variants/codon_variants.csv) that is available in the GitHub repository. Additionally, see https://dms-vep.org/LASV_Josiah_GP_DMS/notebooks/analyze_pacbio_ccs.html for details of CCS quality checking, https://dms-vep.org/LASV_Josiah_GP_DMS/notebooks/build_pacbio_consensus.html for details of creating variant-barcode consensus sequences, and https://dms-vep.org/LASV_Josiah_GP_DMS/notebooks/build_co <u>don_variants.html</u> for details of creating barcode-variant lookup table.

### Short-read Illumina sequencing to read barcodes after selections

Once GPC variants are linked to barcodes using long-read PacBio sequencing (as described in previous section), we can use short-read Illumina sequencing to sequence the variant barcodes to determine mutational frequencies for future experiments. To target non-integrated lentivirus genomes, we trypsinized the cells, washed the cells with PBS, and miniprepped each well using QIAprep Spin Miniprep Kit (Cat. No. 27106X4) approximately 12 hours after infecting the cells.^101,102^ Each miniprepped sample was eluted in 40 µL of EB. Again note, we use non-integrated lentivirus genomes as sequencing templates for sequencing because they are more abundant than the integrated provirus.^103–105^ After cells have been infected and non-integrated lentivirus genomes have been recovered via mini-prepping of the infected mammalian cells, we use a two-step PCR strategy to amplify barcodes for sequencing. The first round of PCR amplifies the barcodes using a forward primer that aligns to the Illumina Truseq Read 1 sequence upstream of the barcode in our lentiviral backbone and a reverse primer that anneals downstream of the barcode and overlaps with the Illumina Truseq Read 2 sequence. The first round PCR was performed using 22 µL of miniprepped sample, 1.5 µL of 10 µM 5’ Illumina round 1 forward primer (IlluminaRnd1_For_CC), 1.5 µL of 10 µM 3’ Illumina round 1 reverse primer (IlluminaRnd1_rev-3_CC), and 25 µL KOD Hot Start Master Mix. Cycling conditions were:

1. 95°C for 2 min
2. 95°C for 20 s
3. 70°C for 1 s
4. 58°C for 10 s, cooling at 0.5°C per 1 s
5. 70°C for 20 s (return to step 2 for another 27x cycles)
6. Hold at 4°C

The PCR products were Ampure XP purified with a sample to bead ratio 1:3. The DNA concentration was measured using a Qubit Fluorometer (ThermoFisher). The second round of PCR uses a forward primer that anneals to the Illumina Truseq Read 1 sequence and has a P5 Illumina adapter overhang, and reverse primers from the PerkinElmer NextFlex DNA Barcode adaptor set that anneals to the Truseq Read 2 site and has the P7 Illumina adapter and i7 sample index. The second round PCR was performed using 20 ng of round one product as determined by Qubit, 2 µL of 10 µM 5’ Illumina round 2 universal forward primer (Rnd2ForUniv_CC), 2 µL of 10 µM 3’ Illumina round 2 indexing reverse primer (Indexing primer), 25 µL KOD Hot Start Master Mix, and remaining volume H2O for a final volume of 50 µL. Cycling conditions were:

1. 95°C for 2 min
2. 95°C for 20 s
3. 70°C for 1 s
4. 58°C for 10 s, cooling at 0.5°C per 1 s
5. 70°C for 20 s (return to step 2 for another 19x cycles)
6. Hold at 4°C

The DNA concentration of the second round PCR products were quantified using a Qubit Fluorometer (ThermoFisher) and all samples were pooled at an even ratio. The pooled samples were gel and Ampure XP purified at a sample to bead ratio of 1:3. The pooled samples were sequenced using either P2 or P3 reagent kits on a NextSeq 1000 or NextSeq2000.

### Selections to determine variant effects on cell entry

To determine the effects of GPC mutations on cell entry as shown in Figure 2, we plated 0.5 million 293T cells per well of a 6-well Poly-L-Lysine tissue culture dish (Corning, Cat. No. 356515). Approximately 24 hours after plating the cells, we infected each well with either ∼4×10^5^ GPC-pseudotyped variant transducing units or ∼1×10^6^ VSV-G-pseudotyped variant transducing units. We aimed to infect cells with at least 8x the number of transducing units as the number of barcoded variants in each library to reduce random bottlenecking of variants. Again note, we use non-integrated lentivirus genomes as sequencing templates because they are more abundant than the integrated provirus, and as a result, transducing units are an underestimate of actual genomes in infected cells.^103–105^ Approximately 12 hours after infecting the cells, we trypsinized the cells, washed the cells with PBS, and extracted non-integrated lentiviral genomes using a QIAprep Spin Miniprep Kit.^101,102^ The variant barcodes were prepared for short-read Illumina sequencing as described in the section above. To determine the effects of GPC mutations on cell entry for cells expressing either human or mastomys DAG1, we followed the same procedure except that we plated 0.5 million 293TΔDAG1+humanDAG1 or 293TΔDAG1+mastomysDAG1 cells per well rather than 293Ts.

To analyze the Illumina short-read sequencing data for variant barcodes, we used the parser (https://jbloomlab.github.io/dms_variants/dms_variants.illuminabarcodeparser.html) implemented in *dms_variants* (https://jbloomlab.github.io/dms_variants). The barcode counts for each variant were converted to scores for the effect on cell entry as previously described in Dadonaite et al.^42^ In brief, scores for the effect on cell entry were calculated for each barcoded variant as log2 (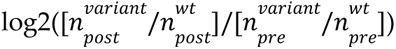) where 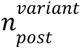 is the count of the variant in the post-selection (GPC-pseudotyped) infection, 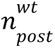 is the count of variants without any mutations (i.e., wildtype GPC) in the post-selection (GPC-pseudotyped) infection, 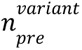 is the count of the variant in the pre-selection (VSV-G-pseudotyped) infection, and 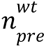 is the count of the variants without any mutations (i.e., wildtype GPC) in the pre-selection (VSV-G-pseudotyped) infection. Positive effects on cell entry meant the variant is better at infecting cells compared to the wildtype GPC while negative effects on cell entry meant the variant is worse at infecting cells compared to the wildtype GPC.

To calculate individual mutation effects on cell entry from the single- and multi-mutant variant scores, we used *multidms* (https://matsengrp.github.io/multidms/)^64^ to fit a global epistasis^70^ model with a sigmoid function. An example global epistasis data fit (https://dms-vep.github.io/LASV_Josiah_GP_DMS/notebooks/func_effects_global_ep <u>istasis_LibA-220823-293T-1.html</u>) can be found as an interactive HTML notebook.

### Production of VSV-G neutralization standard virus

To convert relative barcode counts to absolute neutralization measurements, we spiked in a small amount of a separately produced only-VSV-G-pseudotyped virus pool carrying known barcodes to act as a non-neutralized standard for any selections using antibodies as shown in Figure 3. These viruses were produced as described previously in Dadonaite et al.^42^

### Anti-GPC antibodies

Anti-GPC antibodies 25.10C, 12.1F, 37.7H, 25.6A, 37.2D, and 8.9F first described in Robinson et al.^28^ were produced based on publicly available sequences (https://github.com/dms-vep/LASV_Josiah_GP_DMS/blob/main/non-pipeline <u>_analyses/GPC_antibodies/antibody_sequences.gb</u>).^107^ To produce the antibodies as human IgG1 antibodies, paired VH and VL in a 1:1 ratio were co-transfected transiently into Expi293F cells (ThermoFisher, Cat. No. A14527). The supernatant was harvested 6 days post-transfection, and IgGs were purified with Protein A agarose (Thermo Fisher Scientific, Cat. No. 20333). IgGs were eluted with 100 mM glycine, pH 3 into 1/10th volume 1 M Tris-HCl pH 8.0. IgGs were then buffer exchanged into PBS pH 7.4.

### Selections to determine variant effects on antibody neutralization

To determine the variant effects on antibody neutralization as shown in Figures 3 and 4, we performed antibody selections at the IC_99_, four fold lower than the IC_99_, and four fold greater than the IC_99_ for the six antibodies described in the previous section. We used a spread of antibody concentrations to better fit the biophysical models used to extrapolate individual mutation effects. To perform an antibody selection, we plated 0.5 million 293T cells per well of a 6-well Poly-L-Lysine tissue culture dish (Corning, Cat. No. 356515). Approximately 24 hours after plating the cells, we spiked the VSV-G neutralization standard virus at ∼1% the total GPC-pseudotyped variant library titer. From the combined pool of virus, we incubate ∼4×10^5^ GPC-pseudotyped variant transducing units with antibody at the desired concentration for 1 hour at 37°C. We aimed to infect cells with at least 8x the number transducing units as the number of barcoded variants in each library to reduce random bottlenecking of variants. Again note, we use non-integrated lentivirus genomes as sequencing templates because they are more abundant than the integrated provirus, and as a result, transducing units are an underestimate of actual genomes in infected cells.^103–105^ After the incubation, the virus and antibody mixture was used to infect a well of the plated cells. Approximately 12 hours after infecting the cells, we trypsinized the cells, washed the cells with PBS, and extracted non-integrated lentiviral genomes using a QIAprep Spin Miniprep Kit.^101,102^ The variant barcodes were prepared for short-read Illumina sequencing as described in the section above.

We analyze the short-read Illumina sequencing data from the antibody selections as previously described in Dadonaite et al.^42^ In brief, *polyclonal* (https://jbloomlab.github.io/polyclonal/)^75^ was used to fit neutralization curves and extrapolate individual mutation effects on antibody neutralization from the single- and multi-mutant variants. An example *polyclonal* data fit (https://dms-vep.github.io/LASV_Josiah_GP_DMS/notebooks/fit_escape_antibody_esc <u>ape_LibB-220706-89F-1.html</u>) can be found as an interactive HTML notebook. The per-mutation escape values are roughly proportional to the increase in log fold-change IC_50_ values.

### Validation of GPC variants using lentivirus pseudovirus assays

To validate the effects of specific mutations as shown in Figures 2, 3, and 7, we cloned GPC sequences with desired mutations by performing PCR reactions with partially overlapping desired mutation-containing primers followed by HiFi assembly. The plasmid sequences were verified by Primordium sequencing. To validate the estimated effects of GPC from natural strains, we ordered expression plasmids containing the natural strain GPC from Twist Bioscience. See https://github.com/dms-vep/LASV_Josiah_GP_DMS/tree/main/non-pipeline_analyses/plasmid_maps/GPC_expre <u>ssion_plasmids</u> for all GPC expression plasmids used in this study. Pseudotyped viruses were produced and titrated using the protocols described in Crawford et al.^73^ Note that for the GPC variants cloned to validate the effects on cell entry, we performed unique virus rescues with independent plasmid preparations for each variant (each replicate shown in Figure 2D is a unique plasmid preparation).

To produce the GPC variants, we plated 1 million 293T cells per well of a 6-well tissue culture dish. Approximately, 24 hours after plating the cells, we transfected 1.135 µg of pHAGE6_Luciferase_IRES_ZsGreen lentivirus backbone (2727_pHAGE6-wtCMV-Luc2-BrCr1-ZsGreen-W-1247.gb), 0.525 µg of Gag/Pol helper plasmid (26_HDM_Hgpm2.gb), and 0.34 µg of GPC variant expression plasmid per well using BioT (Bioland Scientific, Cat. No. B01-02) following manufacturer’s protocol and recommendations for reagent ratios. The supernatant was collected 48 hours after transfection and passed through a 0.45 µM SFCA syringe filter.

To titrate GPC variants for validation of effects on cell entry, we plated 20,000 293T cells per well of a clear bottom, poly-L-lysine coated, black walled 96-well plate (Thomas Scientific, Cat. No. 1183B22). Approximately, 24 hours after plating cells, we performed a minimum of two replicate serial dilutions for each rescued pseudotyped virus and infected the plated cells. We then measured luciferase expression at each dilution using Bright-Glo Luciferase Assay System (Promega, E2610) 48 hours after infecting the cells. Virus titers were calculated as relative light units (RLU) per µL for each dilution averaged across dilutions within a linear range.

To perform neutralization assays with the GPC variants, we plated 20,000 293T cells per well of a clear bottom, poly-L-lysine coated, black walled 96-well plate (Thomas Scientific, Cat. No. 1183B22). Approximately, 24 hours after plating cells, we performed replicate serial dilutions for each antibody and then incubated each antibody dilution with GPC variant virus for one hour. After the incubation, we infected the plated cells. We then measured luciferase expression at each dilution using Bright-Glo Luciferase Assay System (Promega, E2610) 48 hours after infecting the cells. Fraction infectivity for each antibody dilution was calculated by subtracting background RLU readings from uninfected cells and dividing RLU readings by RLU readings for cells infected without any antibody. Fraction infectivities were used to fit neutralization curves using *neutcurve* (https://jbloomlab.github.io/neutcurve/).

### Analysis of effects of mutations on cell entry for human and mastomys ɑ-DG-mediated entry

To determine if there were any biologically significant differences between cell entry mediated by human or mastomys *ɑ*-DG as shown in Figure S5, we used *multidms* (https://matsengrp.github.io/multidms/).^64^ In brief, *multidms* applies a custom global epistasis model^70^ to calculate shifts from the experimental condition (i.e., selections conducted on 293TΔDAG1+humanDAG1 or 293TΔDAG1+mastomysDAG1 cells) to a reference condition (i.e., selections conducted on 293T cells). To determine what shifts are biologically significant, we apply a lasso regularization sweep with different lasso penalty weights that encourage the shift values to be zero (Figure S5B-D). We picked a lasso weight where the majority of stop codons converged to zero because it would be expected that stop codons would be equally deleterious for entry into cells regardless of the host receptor present. Both missense and nonsense mutation shift values converged to zero at similar rates supporting that there is not a detectable difference between the *ɑ*-DG orthologs (Figure S5D). The computational script used to analyze the mutational shift values for the different conditions (https://dms-vep.github.io/LASV_Josiah_GP_DMS/notebooks/human_mastomys_correlation <u>.html</u>) is included as an html notebook.

### DAG1 alignment and sequence analysis

To create a list of *DAG1* sequences, all protein sequences for “DAG1 - dystroglycan 1” as of 29 September 2023 were downloaded from NCBI. For the human and mastomys *DAG1* alignment in Figure S2, we extracted the *DAG1* sequences for human and mastomys and aligned them using MAFFT.^108^ In addition, we aligned all *DAG1* sequences using MAFFT^108^ to analyze the conservation across species at the glycosylation motif required for Lassa GPC attachment as shown in Figure S2.^61^ Note that the full length mastomys *DAG1* sequence is from *Mastomys coucha* because this is the only fully annotated *DAG1* sequence available for the *Mastomys* genus. It has previously been shown that *ɑ*-DG is 100% identical at the amino-acid level between *Mastomys natalensis* and *Mastomys coucha*^95^ and only *ɑ*-DG affects Lassa virus GPC attachment (not β-DG).^96^ The computational pipeline used for this analysis (https://github.com/dms-vep/LASV_Josiah_GP_DMS/tree/main/non-pipeline_analyses/DAG1_phylogeny_analysis) is included as a subdirectory in the GitHub repo.

### Natural GPC sequence analysis

To create a curated list of GPC sequences, all accessions for “Mammarenavirus lassaense (taxid:3052310)” as of 10 August 2023 were downloaded from NCBI Virus. GPC sequences were extracted from the sequences using EMBOSS tools.^109^ Only sequences with full length GPC sequences and no ‘N’ characters were retained. For the reduced phylogeny in Figure 1C, a subset of representative GPC sequences were selected using CD-HIT.^110^ We then aligned the representative GPC sequences using MAFFT.^108^ Codon alignments were created from the protein alignments using a custom Python script (https://github.com/dms-vep/LASV_Josiah_GP_DMS/blob/main/non-pipeline_analyses/ <u>LASV_phylogeny_analysis/Scripts/create_codon_alignment.py</u>). A phylogenetic tree was inferred from the codon alignment using IQ-TREE^111^ and rendered using ETE3^112^ as shown in Figure 1C.

To calculate site variability of the GPC as shown in Figure S6D,E we aligned all extracted GPC protein sequences using MAFFT^108^ and then calculated the effective number of amino acids (exponential of Shannon entropy) per site.^94^ Amino-acid mutation frequencies relative to the Josiah strain were also calculated from the protein multiple sequence alignment. The computational pipeline used to analyze all Lassa GPC sequences and construct phylogenetic trees (https://github.com/dms-vep/LASV_Josiah_GP_DMS/tree/main/non-pipeline_analyses/LASV_ph <u>ylogeny_analysis</u>) is included as a subdirectory in the GitHub repo.

### Short-read sequencing data analysis for LM395 strain

The N89D mutation present in the LM395 strain was reported to be a high-frequency polymorphism in Andersen et al^17^ but we wanted to independently verify this by reanalyzing the short-read sequencing data to determine the base calling at position 89. This analysis was conducted using an automated computational pipeline (https://github.com/dms-vep/LASV_Josiah_GP_DMS/tree/main/non-pipeline_analyses/LASV_NGS_analysis) that is included as a subdirectory in the GitHub repo. In brief, we downloaded the sequencing run information for the LM395 strain from BioProject: PRJNA254017. The raw FASTQ files were first trimmed for adapter sequences and trimmed of homopolymer sequences (>10) at the end of the reads as well as filtered for reads based on a minimum Phred score (<25) and minimum length (<25bp) using fastp.^113^ Duplicate reads were removed using SAMtools.^114^ Reads were mapped to the Josiah Lassa reference strain (S segment: NC_004296 and L segment: NC_004297) using BWA mem.^115^ Aligned data was summarized using SAMtools^114^ and parsed using a custom Python script (https://github.com/dms-vep/LASV_Josiah_GP_DMS/blob/main/non-pipeline_analyses/LASV_NGS_analysis/Scripts/ <u>process_mpileup_file.py</u>).

### Structural analysis

To provide structural context for the deep mutational scanning data as shown in Figures 2B, 4B, 5, S7B-D, and S8B, mutational effects were mapped onto the surface representation of the full-length native GPC structure (PDB: 7PUY^52^) using PyMol. To visualize antibody structures with the antibody escape maps, antibody structures (25.10C PDB: 7TYV,^51^ 12.1F PDB: 7UOV,^31^ 37.7H PDB: 5VK2,^48^ 25.6A PDB: 6P95,^49^ 37.2D PDB: 7UOT,^31^ and 8.9F PDB: 7UOT^31^) were aligned to the full-length native GPC structure (PDB: 7PUY^52^) using PyMol. Antibody contacts were determined as any non-hydrogen GPC atoms within 4 Å of the antibody using Bio3d.^116^ The full computational script used to calculate antibody contacts (https://github.com/dms-vep/LASV_Josiah_GP_DMS/tree/main/non-pipeline_analyses/pd <u>b_antibody_contacts</u>) is included as a subdirectory in the GitHub repository.

GPC regions shown in Figures 2, S6, and S7 were identified as the following: stable signal peptide (SSP) represents sites 1 to 58, glycoprotein 1 (GP1) represents sites 59 to 259, glycoprotein 2 (GP2) represents sites 260 to 491, transmembrane domain (TM) represents sites 428 to 447,^93^ C-terminal cytoplasmic tail (CT) represents sites 448 to 491,^93^ *ɑ*-DG binding sites represent sites described in Katz et al^52^ that are involved in *ɑ*-DG attachment, LAMP1 binding sites represent sites described in Cohen-Dvashi et al^72^ and Israeli et al^71^ that are important for LAMP1 attachment, and N-glycosylation sites represent the 11 N-linked glycosylation motif sites (N-X-S/T, X ≠ P) present in the Josiah strain GPC at sites N79, N89, N99, N109, N119, N167, N224, N365, N373, N390, and N395.

## Resource availability

### Lead contact

Further information and requests for reagents and resources should be directed to and will be fulfilled by the lead contact, Jesse Bloom.

### Data and Code Availability

All computer code and data are available at https://github.com/dms-vep/LASV_Josiah_GP_DMS and the raw sequencing data for this study can be found in the NCBI Sequence Read Archive under BioProject PRJNA1071644.

### Materials Availability

A lentiviral backbone appropriate for the deep mutational scanning approach used here and the corresponding helper plasmids are available in AddGene under item numbers 204146, 204152, 204153, and 204154. Lassa GPC mutant libraries generated in this study will be made available on request by the lead contact with a completed Materials Transfer Agreement.

---

**Figure S1.**
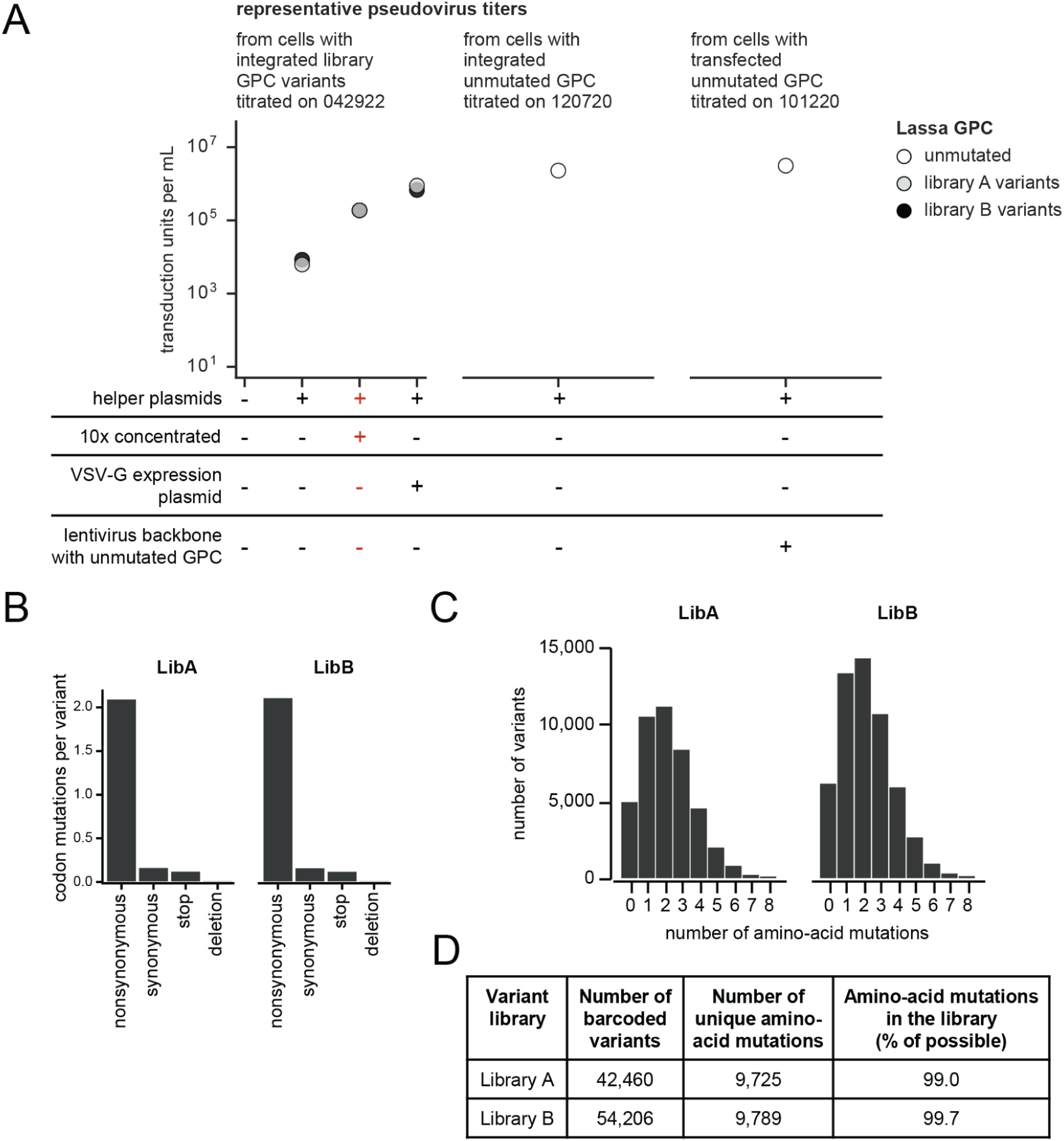
Characteristics of GPC deep mutational scanning libraries, related to Figure 1. **A** GPC-pseudotyped lentivirus titers. Pseudotyped viruses were generated from 293T-rtTA cells with a single GPC variant integrated in each cell as depicted in Figure 1B or from transfection of lentivirus backbone containing GPC. Viruses were produced using the indicated conditions and titrated on 293T cells. The GPC-pseudotyped variant viruses used for deep mutational scanning were generated under the conditions highlighted in red. The VSV-G condition was used to generate VSV-G-pseudotyped variant viruses, which were used to assess the composition of mutations present in the variant libraries. **B** Average number of codon mutations per barcoded variant for each GPC-pseudotyped variant library. **C** Distribution of the number of amino-acid mutations in each GPC variant for each library. **D** The total number of barcoded variants, unique amino-acid mutations, and percentage of possible amino-acid mutations for each GPC-pseudotyped variant library.

**Figure S2.**
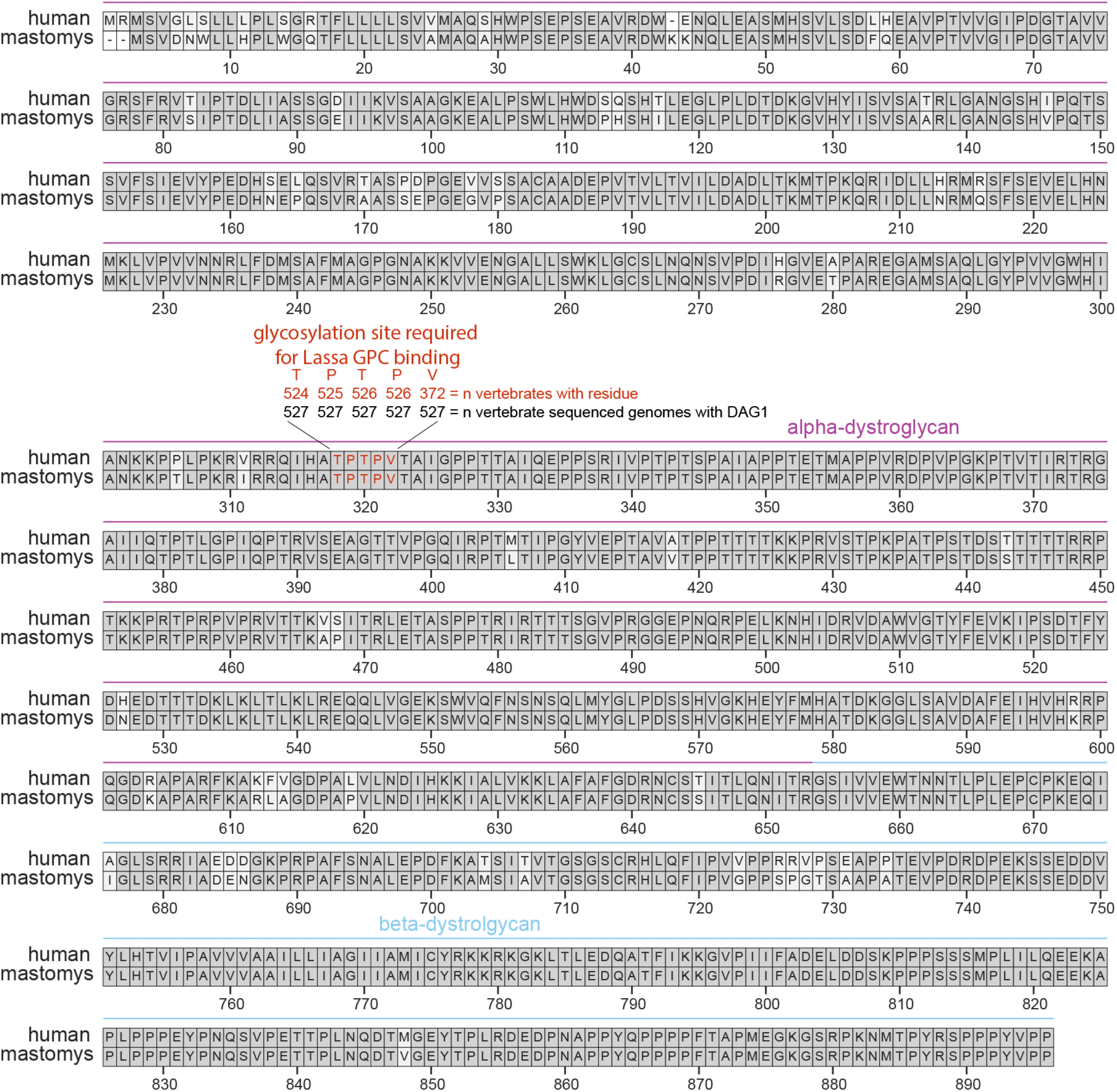
Human and mastomys DAG1 sequence alignment, related to Figure 2. Protein sequence alignment of human and mastomys DAG1. The alpha-dystroglycan (*ɑ*-DG) and beta-dystroglycan (β-DG) subunits are highlighted in purple and blue, respectively. The TPTPV glycosylation motif required for laminin and Lassa virus GPC binding^61^ is highlighted in red with the conservation across all vertebrate sequences shown in the text. The overall conservation between human and mastomys *ɑ*-DG at the protein level is 93%.

**Figure S3.**
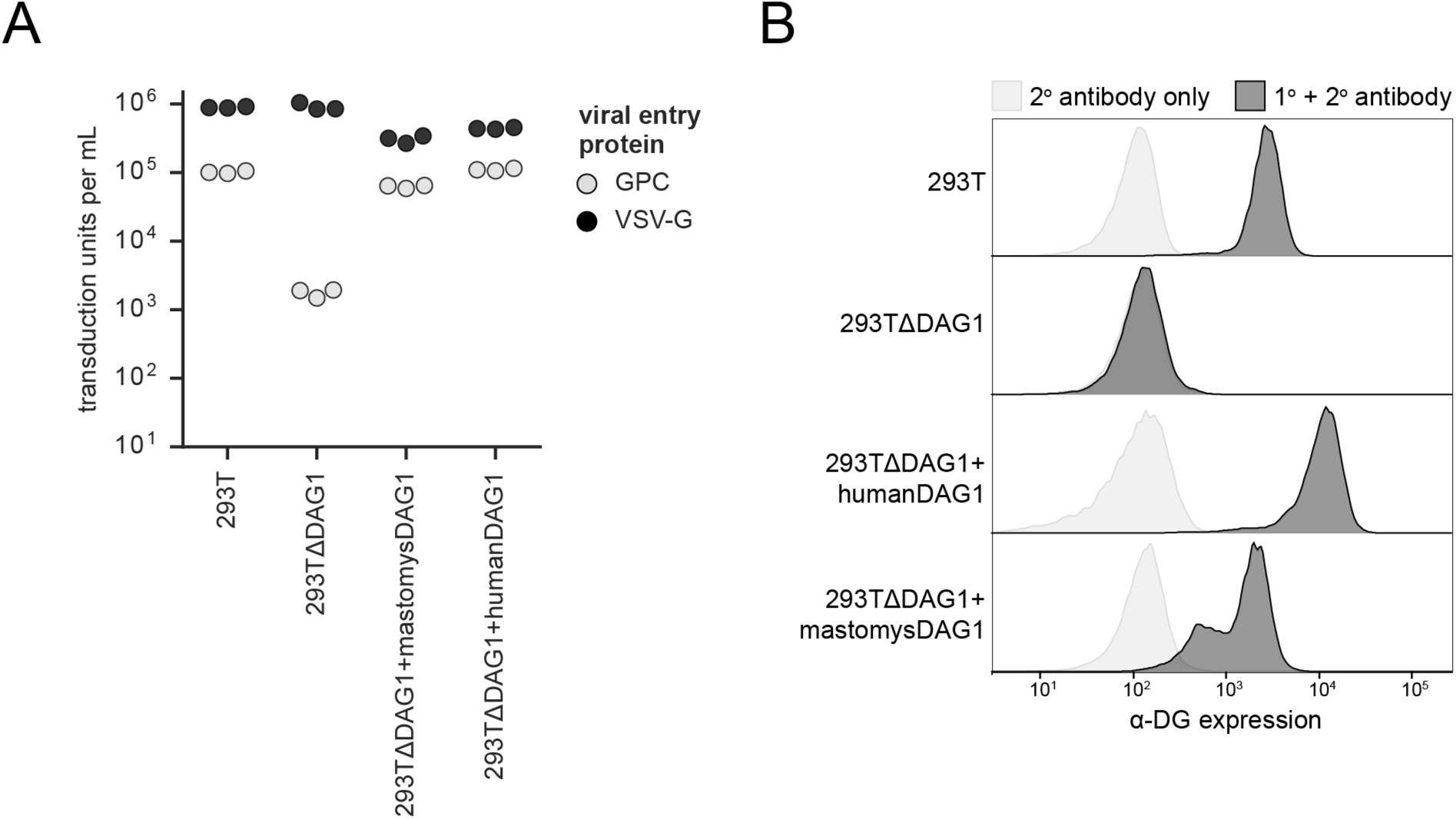
GPC mediates effective entry into cells expressing human or mastomys DAG1 but not cells lacking DAG1, related to Figure 2. **A** Titers of VSV-G or Lassa GPC pseudotyped lentivirus in different cell lines. 293T cells natively express human *ɑ*-DG from their *DAG1* gene, 293TΔDAG1 have had this gene disrupted and so do not express *ɑ*-DG,^27^ and 293TΔDAG1+mastomysDAG1 / 293TΔDAG1+humanDAG1 cells are variants of 293TΔDAG1 that we stably transduced with the mastomys or human *DAG1* gene. VSV-G-mediated cell entry is independent of *DAG1* expression since VSV-G uses a different receptor, but GPC-mediated cell entry is only efficient in the presence of *ɑ*-DG expression from the *DAG1* gene. **B** *ɑ*-DG expression for different cell lines. Surface *ɑ*-DG expression was measured using flow cytometry with an antibody against dystroglycan and the histograms show the distribution of expression over a population of cells. See “Methods” for details.

**Figure S4.**
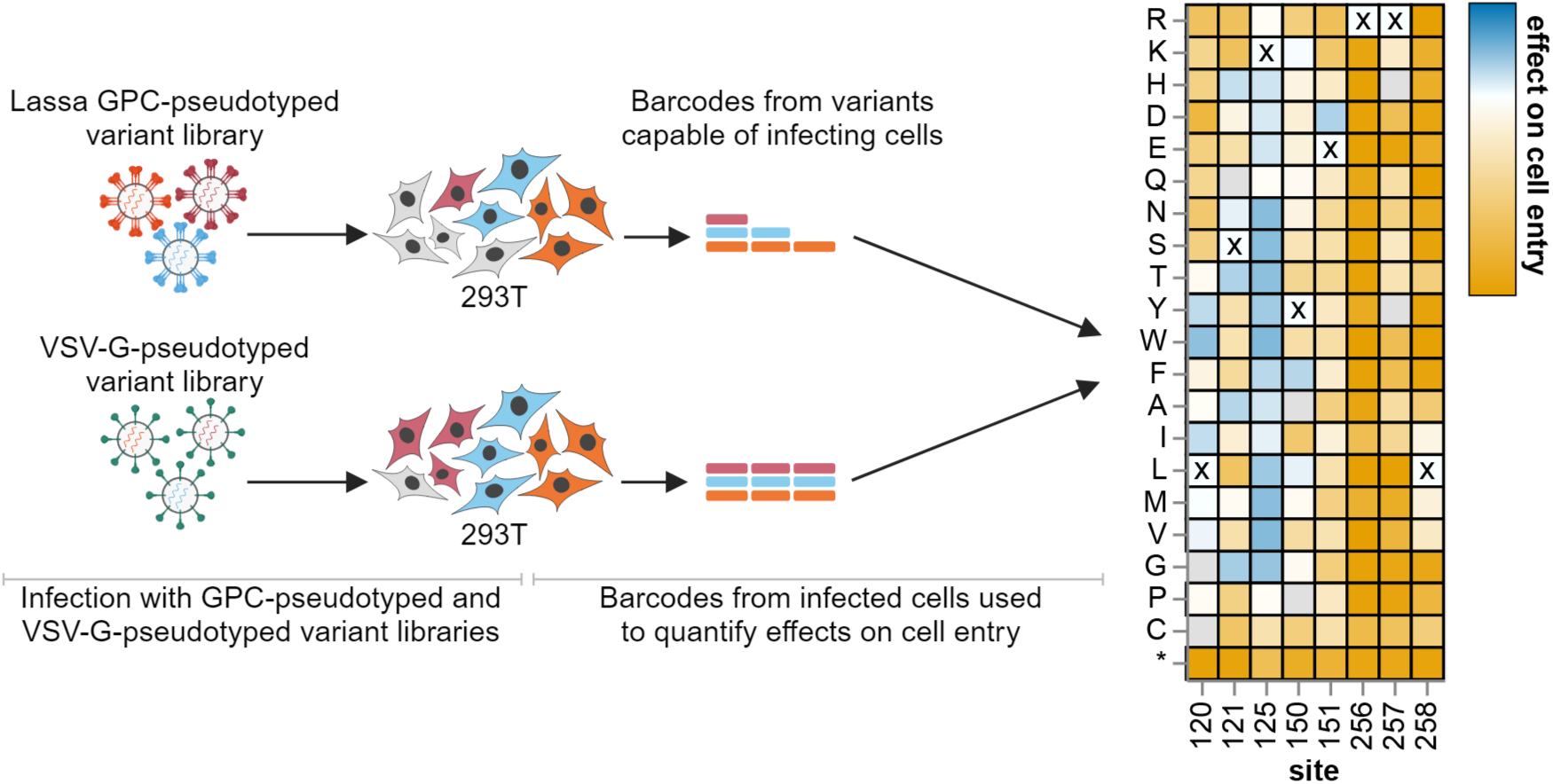
Workflow for measuring how mutations affect cell entry, related to Figure 2. Cells are infected with either the GPC-pseudotyped variant library or the VSV-G-pseudotyped library, and variant barcodes are sequenced from the infected cells. All variants infect cells when VSV-G is present, but only variants with functional GPC infect cells when VSV-G is not present. The log enrichment or depletion of variants in the GPC-pseudotyped condition relative to the VSV-G-pseudotyped condition quantifies the functional capability of variants to enter cells. Individual mutation effects are inferred from the single- and multi-mutant variants using a global epistasis model.^70^

**Figure S5.**
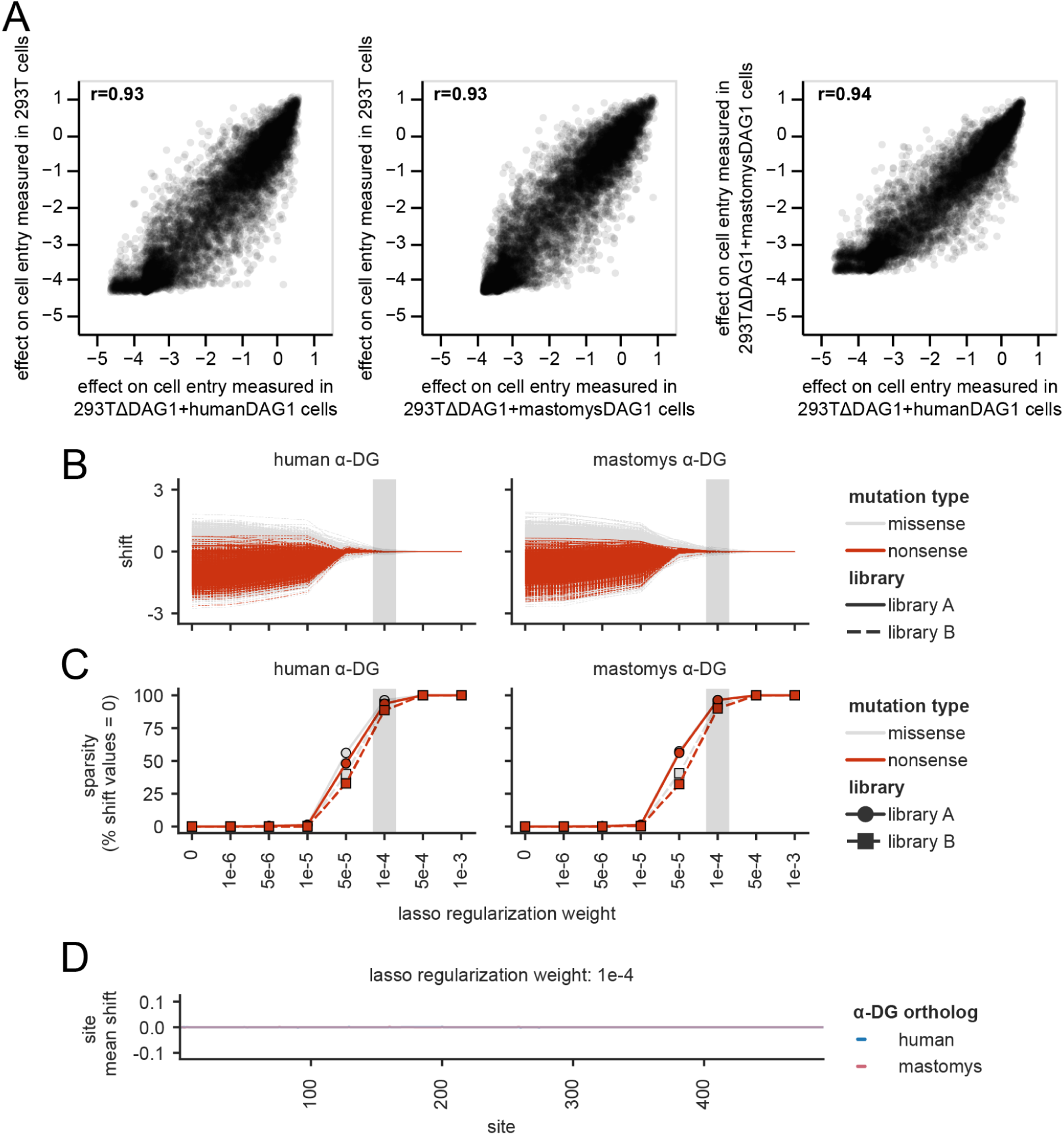
Effects of GPC mutations on entry into 293T cells expressing human or mastomys ɑ-DG, related to Figure 2. **A** Correlation of mutation effects on cell entry measured in 293T, 293TΔDAG1+humanDAG1, and 293TΔDAG1+mastomysDAG1 cells. See https://dms-vep.org/LASV_Josiah_GP_DMS/htmls/DAG1_ortholog_correlations.html for an interactive version of correlation plots that allow you to mouse over points for amino-acid identities. The Pearson correlation (r) is indicated. For interactive plots of the effects of all mutations on entry, see the following links: for 293T cells (https://dms-vep.org/LASV_Josiah_GP_DMS/htmls/293T_entry_func_effects.html), for 293TΔDAG1+humanDAG1 (https://dms-vep.org/LASV_Josiah_GP_DMS/htmls/human_293T_ <u>entry_func_effects.html</u>), and for 293TΔDAG1+mastomysDAG1 (https://dms-vep.org/LASV_Josiah_GP_DMS/htmls/mastomys_293T_entry_func_effects.html). **B-D** Analysis of possible shifts in mutation effects in cells expressing human versus mastomys *ɑ*-DG as assessed using the more sophisticated algorithm implemented in the *multidms* software package (https://github.com/matsengrp/multidms).^64^ **B** Shifts in mutation effects on cell entry measured on 293TΔDAG1+human and 293TΔDAG1+mastomys cell lines relative to 293T cells. Each line tracks the shift value for a single mutation across different lasso weight regularization values. Red lines are for all nonsense mutations and gray lines are for all missense mutations. Individual library measurements are shown in solid and dashed lines. **C** The sparsity (i.e., percent shift values equal to zero) across different lasso weight regularization values. Lines are colored as indicated in **B**. **D** Per site average shift value for mutations measured on 293TΔDAG1+human and 293TΔDAG1+mastomys cell lines for a lasso weight of 1e-4. This is a reasonable lasso weight because it regularizes apparent shifts for stop codons (which should not have human versus mastomys specific effects) to zero. At this lasso weight, the flat line in this plot shows that there are no sites with an appreciable shift in mutation effects.

**Figure S6.**
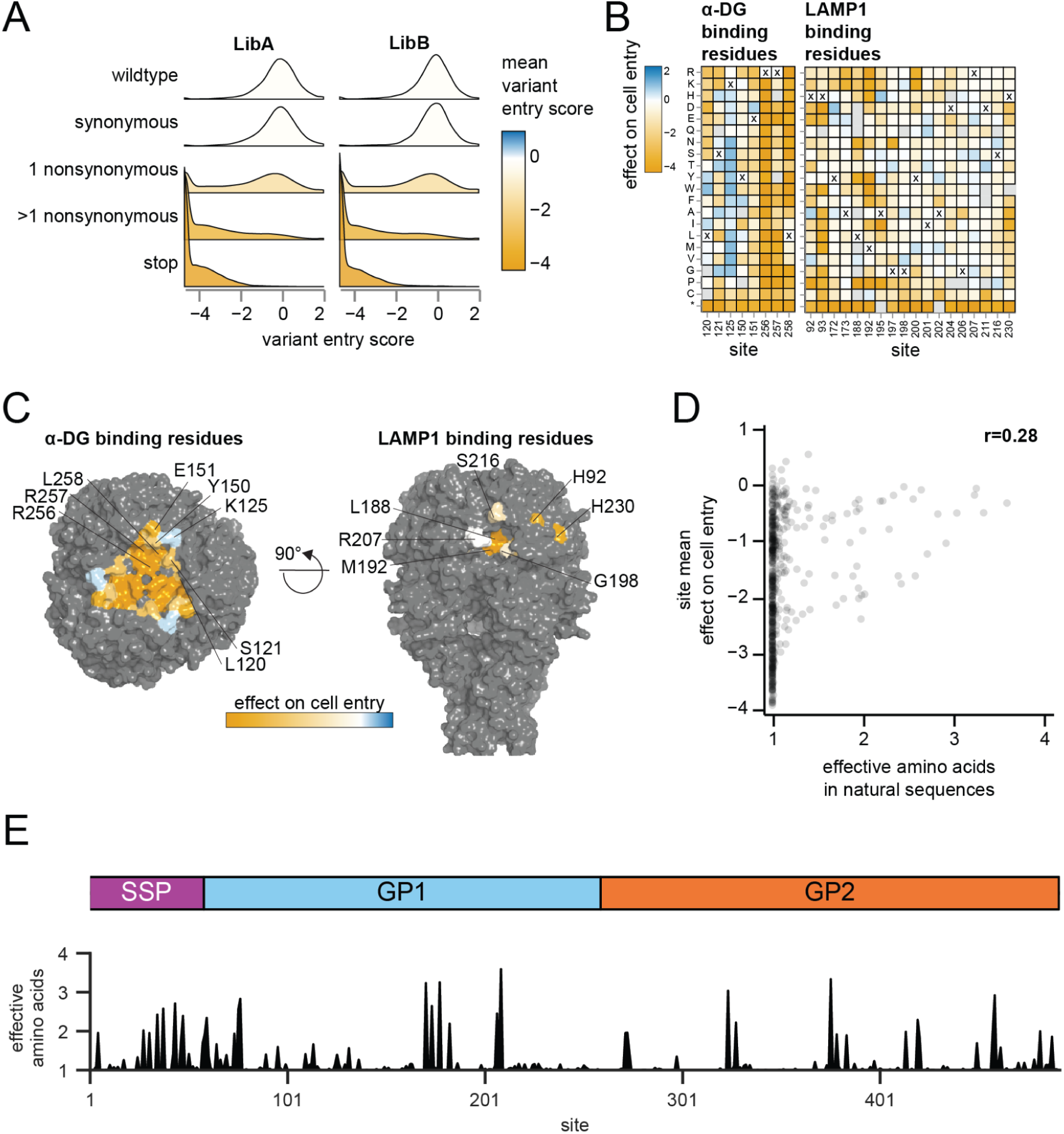
Effects of mutations on cell entry for specific GPC regions and in comparison to natural sequence diversity, related to Figure 2. **A** Distributions of deep mutational scanning cell-entry scores for variants with different mutant types. The cell-entry score is the log ratio of the variant frequency relative to the unmutated parental strain in the GPC-pseudotyped library versus the VSV-G-pseudotyped library. Negative and positive scores indicate decreased and increased GPC-mediated cell entry relative to unmutated GPC, respectively. Distributions are colored by mean cell-entry score. **B** Heatmaps of effects on cell entry of individual mutations at key sites for host receptor binding. Sites involved in *ɑ*-DG^52^ binding and sites suggested to be involved in LAMP1^71,72^ binding are shown. **C** Surface representation of Lassa GPC colored by per site average amino-acid effect on cell entry for key sites for *ɑ*-DG and LAMP1 binding (PDB: 7PUY). **D** Correlation of per site average effects of amino-acid mutations on cell entry measured by deep mutational scanning and natural sequence diversity as described in **E**. **E** Top shows a schematic of Lassa GPC with the stable signal peptide (SSP), glycoprotein 1 (GP1), and glycoprotein 2 (GP2) highlighted. Below is a plot of natural sequence diversity across an alignment of all high quality Lassa GPC sequences. Diversity is quantified as the effective number of amino acids at a site.^94^

**Figure S7.**
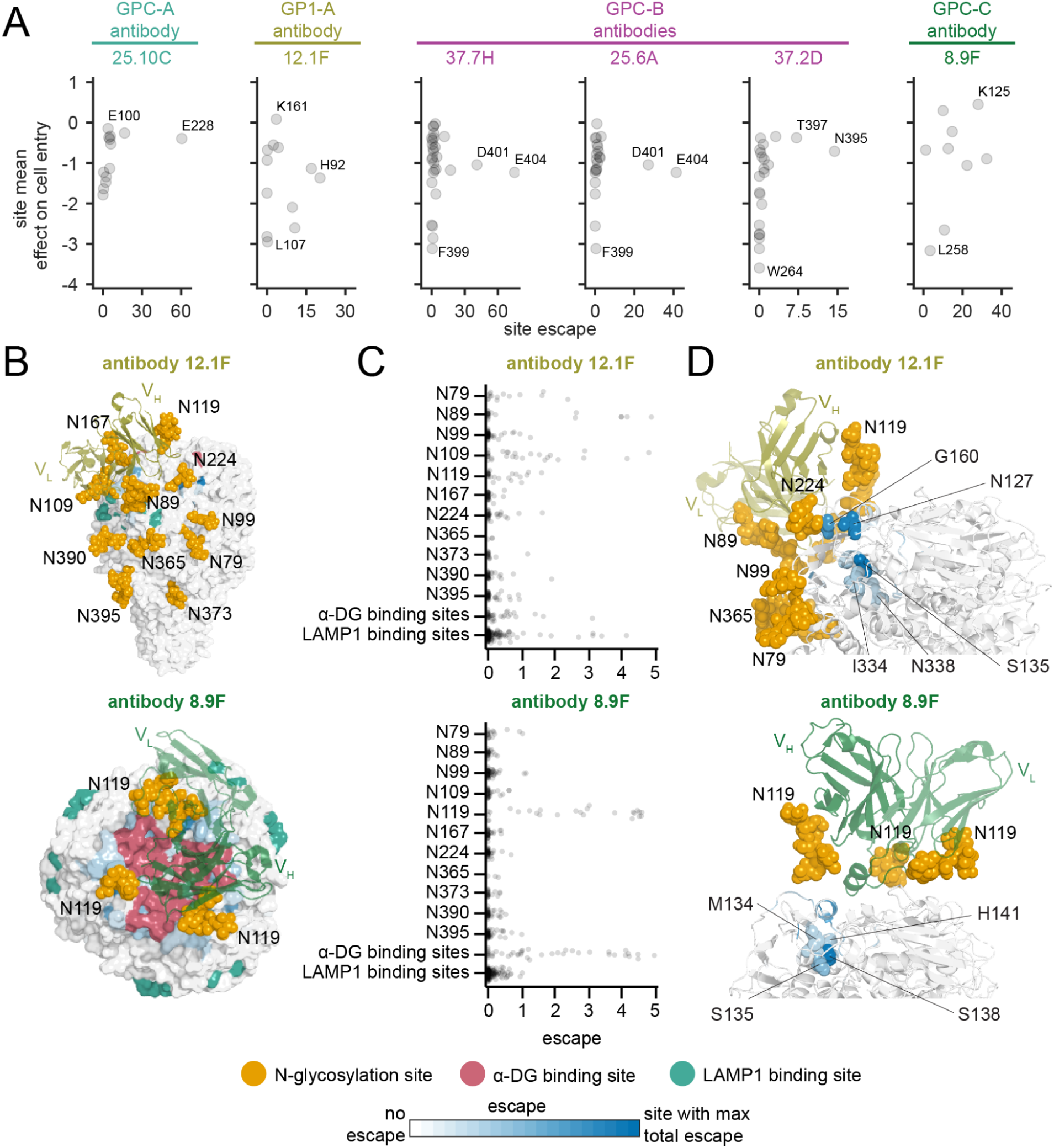
Effects of mutations on antibody escape in comparison to effects on cell entry and for specific GPC sites, related to Figures 4 and 5. **A** Comparison of per site average effects of amino-acid mutations on cell entry and per site summed effects of mutations on antibody escape for all sites that contact the antibody (within 4 Å). For instance, site L258 (which contacts antibody 8.9F) has low mutational tolerance, which may explain why it is not measured to cause escape as our assay can only quantify escape for mutations tolerated for GPC-mediated cell entry. On the other hand, site E100 (which contacts antibody 25.10C) has low escape even though mutations there are well tolerated, suggesting it is simply not important for antibody binding. The antibody escape maps are grouped by antibody epitope classification of Robinson et al.^28^ See https://dms-vep.org/LASV_Josiah_GP_DMS/htmls <u>/antibody_escape_vs_func_effect.html</u> for an interactive version of this plot. **B** Surface representation of Fab-bound GPC colored by site escape as measured in deep mutational scanning, with the Fab shown in a colored cartoon representation. Antibody 12.1F inhibits receptor binding and relies on five N-linked glycans (N89, N109, N119, N167, and N224) to bind to a single monomer of GPC^31,78^; therefore, mutations to glycosylation motifs (N-X-S/T, X≠P) and to receptor binding sites escape antibody 12.1F neutralization as shown in **C**. Similarly, antibody 8.9F inhibits *ɑ*-DG binding and relies on the N-linked glycan N119 to bind across the three monomers of GPC^31^; therefore, mutations to *ɑ*-DG binding sites and to the N119 glycosylation motif escape 8.9F neutralization as shown in **C**. Blue indicates the GPC site with the most escape from that antibody, and white indicates sites with no escape. N-linked glycans (orange spheres), *ɑ*-DG binding sites (pink), and LAMP1 binding sites (turquoise) are highlighted. The Fab bound antibody structures shown here come from prior cryo-EM structures.^31^ **C** Effects of mutations on antibody escape for N-linked glycosylation sites (N-X-S/T, X≠P) and receptor binding sites. Mutations affecting glycans that are important for antibody binding (e.g., antibody 8.9F relies on N-linked glycan N119) tend to lead to antibody escape. Each point represents a different amino-acid mutation. **D** Zoomed in view of cartoon representation of Fab-bound GPC colored by site escape as measured in deep mutational scanning, with the Fab shown in a colored cartoon. Antibody escape sites that are more distally located from the antibody are highlighted by spheres. For instance, site S135 is neither an antibody 12.1F contact nor part of a glycosylation motif that is important for 12.1F binding, but mutations to this site lead to antibody escape.

**Figure S8.**
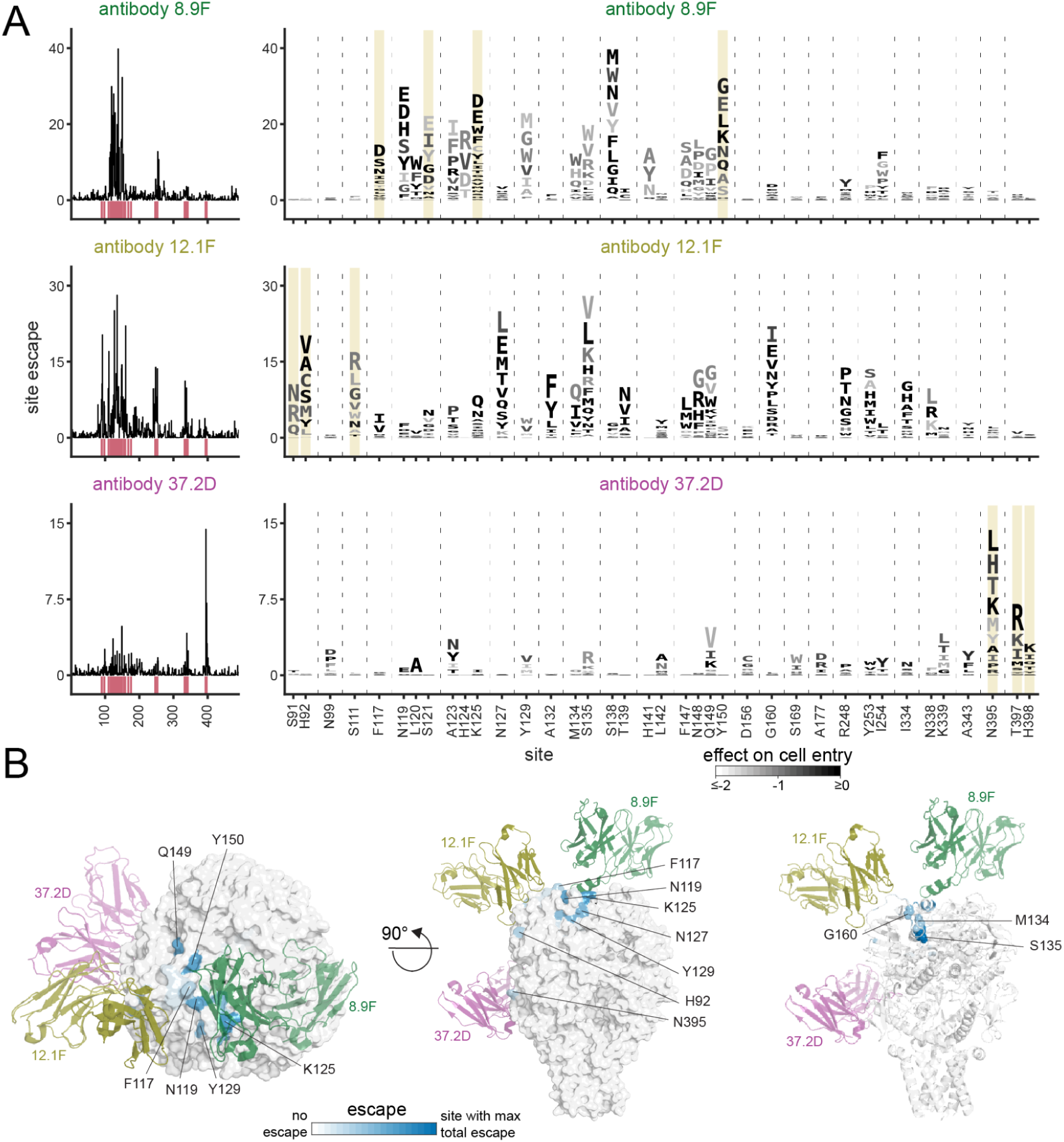
Effects of mutations on antibody escape for the antibodies that comprise the antibody cocktail Arevirumab-3, related to Figures 4 and 5. **A** Escape maps for the antibodies (8.9F, 12.1F, and 37.2D) that comprise the antibody cocktail Arevirumab-3. Line plots show site summed effects of all escape mutations at a site as measured in the deep mutational scanning. The top 15 escape sites across the three antibodies are highlighted pink below the line plot and shown in logo plots where the height of each letter indicates escape caused by that mutation. Letters are colored by mutational effects on cell entry in the absence of antibody, with mutations that impair entry shown in lighter gray. Sites that contact the antibody (within 4 Å) are highlighted in yellow. **B** Surface representation of Fab-bound GPC colored by site escape averaged across the three Arevirumab-3 antibodies (8.9F, 12.1F, and 37.2D) as measured in deep mutational scanning, with the Fab of the antibodies shown in colored cartoon representations. Because GPC is a homo-trimer, escape is colored only on sites in the monomer that is closest to the antibodies shown. Blue indicates the GPC site with the most escape from that antibody, and white indicates sites with no escape. The Fab bound antibody structures shown here come from prior cryo-EM structures.^31^ See beginning of “Methods” for links to more detailed interactive versions of each escape map.

**Figure S9.**
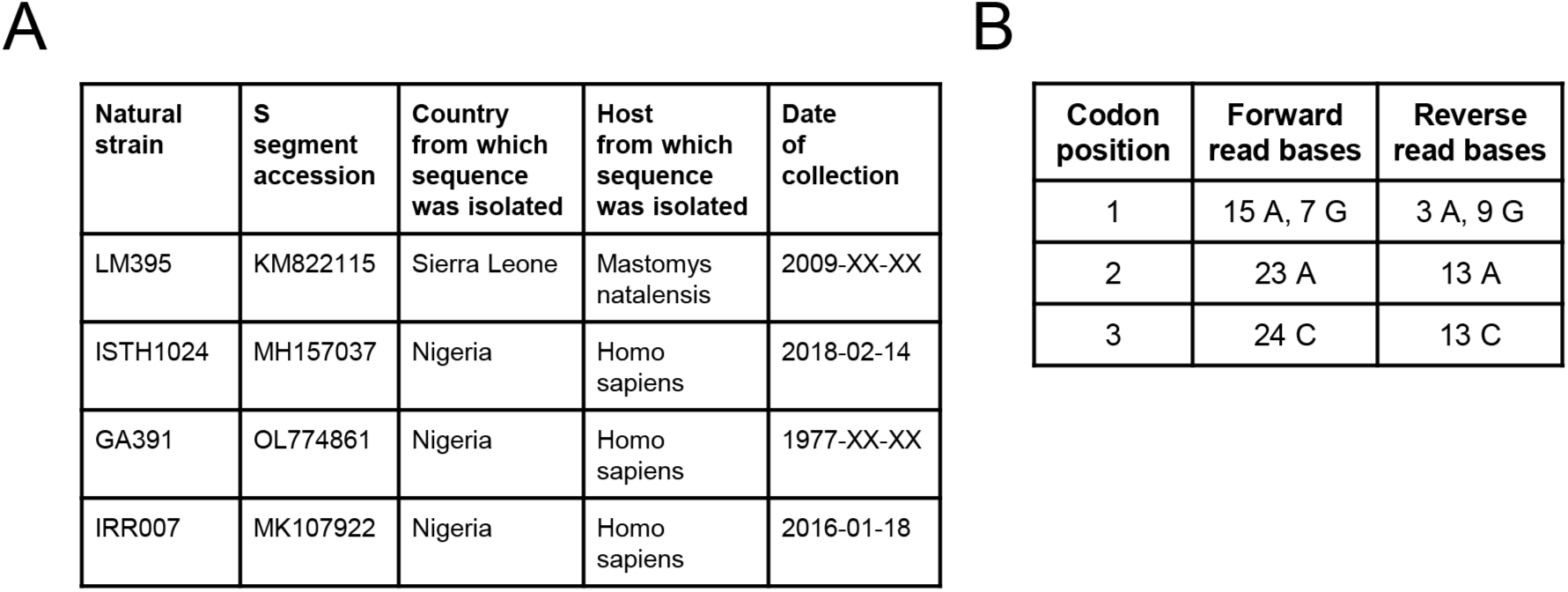
Natural strain GPCs used for neutralization assays, related to Figures 6 and 7. **A** Additional metadata for natural strain GPCs used for experiments. **B** Short-read sequencing data for LM395 strain at codon position 89. The columns indicate read counts in the forward and reverse strands. The AAC codon encodes N (the identity in Josiah) and the GAC codon encodes D (the polymorphic mutant identity in LM395 strain at position 89). See “Methods” for more details on analysis.

**Figure S10.**
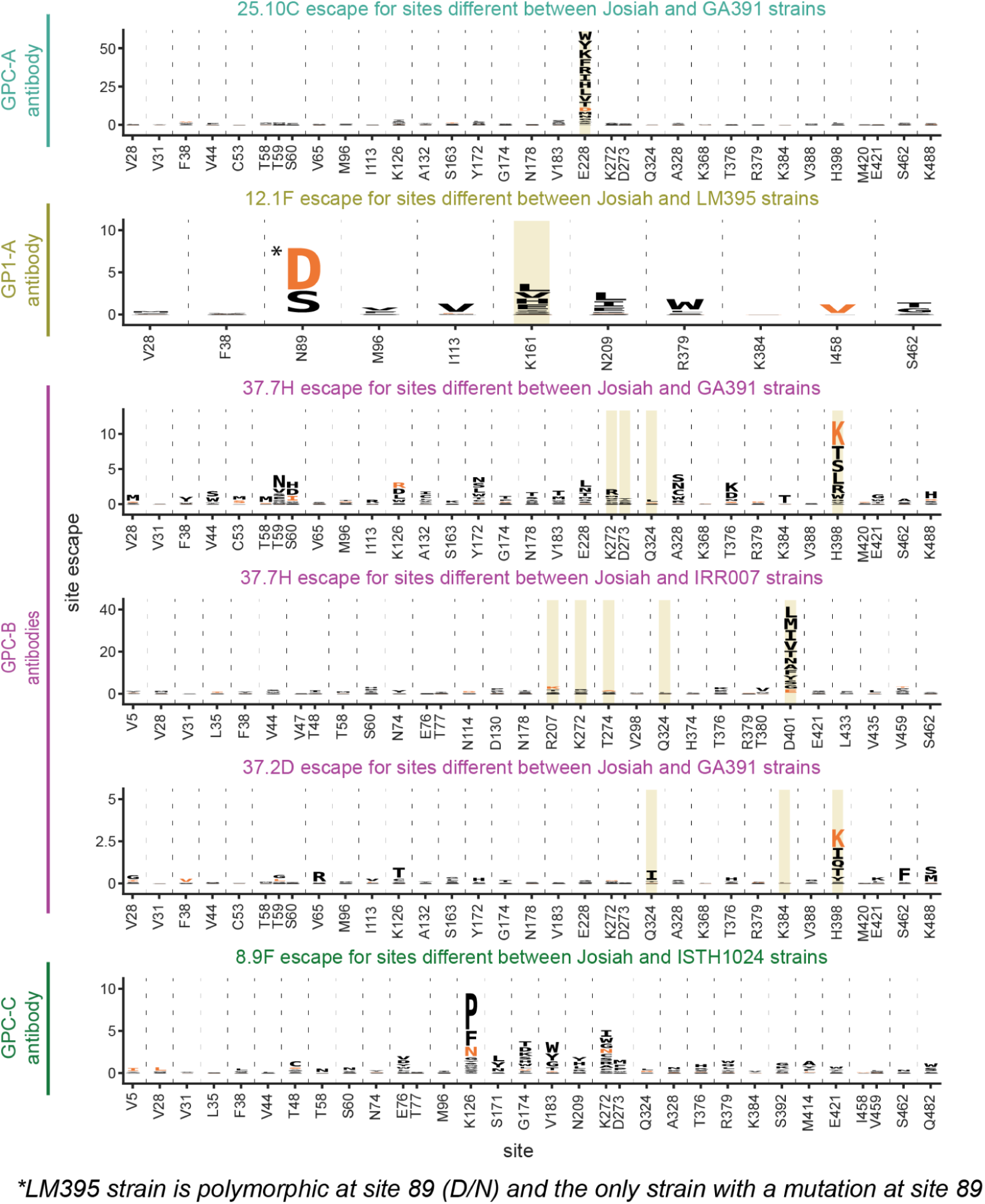
Escape maps for all sites that differ between Josiah and natural strain GPCs, related to Figure 6 and 7. Antibody escape maps for all amino-acid sites that differ between the Josiah GPC strain and the natural strain GPC chosen for validation. This plot differs from Figure 7A as it shows all sites that differ between the GPCs, rather than just the top 10 escape sites. As in Figure 7A, the height of the letter corresponds to the strength of escape, and the amino acid present in the natural isolate is colored orange. Sites that contact the antibody (within 4 Å of antibody) are highlighted yellow. The antibody escape maps are grouped by antibody epitope classification of Robinson et al.^28^ *N89D is marked because the LM395 strain with the N89D mutation is polymorphic at site 89 (Figure S9B)^17^ and is the only strain with a mutation at site 89.

